# *Tetrahymena* RIB72A and RIB72B are Microtubule Inner Proteins in the ciliary doublet microtubules

**DOI:** 10.1101/356428

**Authors:** Daniel Stoddard, Ying Zhao, Brian A. Bayless, Long Gui, Panagiota Louka, Drashti Dave, Swati Suryawanshi, Raphaël F.-X. Tomasi, Pascale Dupuis-Williams, Charles N. Baroud, Jacek Gaertig, Mark Winey, Daniela Nicastro

**Affiliations:** Department of Biology, Rosenstiel Basic Medical Sciences Research Center, Brandeis University, Waltham, MA 02453, USA; Departments of Cell Biology and Biophysics, University of Texas Southwestern Medical Center, Dallas, TX 75390, USA; Department of Molecular, Cellular & Developmental Biology University of Colorado Boulder, Boulder, Colorado 80309, USA; Department of Molecular and Cellular Biology, University of California, Davis, CA, 95616, USA; Department of Cellular Biology, University of Georgia, Athens, GA, 30602, USA; Department of Mechanics, LadHyX, CNRS and Ecole Polytechnique, 91128 Palaiseau Cedex, France; UMR-S 1174 Inserm, Universite Paris-Sud, Bat 443, 91405 Orsay Cedex, France; ESPCI, Paris France

## Abstract

Doublet and triplet microtubules are essential and highly stable core structures of centrioles, basal bodies, cilia and flagella. In contrast to dynamic cytoplasmic microtubules, their luminal surface is coated with regularly arranged Microtubule Inner Proteins (MIPs). However, the protein composition and biological function(s) of MIPs remain poorly understood. Using genetic, biochemical and imaging techniques we identified *Tetrahymena* RIB72A and RIB72B proteins as ciliary MIPs. Fluorescence imaging of tagged RIB72A and RIB72B showed that both proteins co-localize to *Tetrahymena* cilia and basal bodies, but assemble independently. Cryo-electron tomography of *RIB72A* and/or *RIB72B* knockout strains revealed major structural defects in the ciliary A-tubule involving MIP1, MIP4 and MIP6 structures. The defects of individual mutants were complementary in the double mutant. All mutants had reduced swimming speed and ciliary beat frequencies, and high-speed video imaging revealed abnormal highly curved cilia during power stroke. Our results show that RIB72A and RIB72B are crucial for the structural assembly of ciliary A-tubule MIPs and are important for proper ciliary motility.

**SUMMARY:** Microtubule Inner Proteins (MIPs) bind to the luminal surface of highly stable microtubules. Combining cell biology and cryo-electron tomography, Stoddard *et al.* show that RIB72A and RIB72B are conserved MIPs in ciliary doublet microtubules and that they are important for proper ciliary motility.

## INTRODUCTION

Centrioles, basal bodies, cilia and flagella are microtubule assemblies that are critical for fundamental cellular functions, including centrosome biogenesis, assembly and organization of the microtubule cytoskeleton, cell sensing and signaling by primary cilia, and propelling cells or generating fluid flow across tissues (Satir, 2017). Defects in ciliary assembly or function lead to a diverse array of human diseases, such as developmental disorders, numerous types of cancer, and ciliary diseases collectively known as ciliopathies. The latter can cause birth defects that affect development of the brain (microcephaly), skeleton and heart (e.g. *situs inversus*), polycystic kidney disease, Bardet-Biedl syndrome, and primary ciliary dyskinesia (Fliegauf *et al.*, 2007; Satir, 2017).

The core structure of cilia/flagella and basal bodies is a highly conserved, cylindrical arrangement of nine doublet or triplet microtubules, respectively. Whereas singlet cytoplasmic microtubules are usually characterized by dynamic instability, doublet and triplet microtubules form a highly stable scaffold, which serves as a platform for a plethora of associated proteins. A common structural feature shared by doublets, triplets, and other highly stable microtubules, such as the subpellicular and ventral disc microtubules of the pathogens *Plasmodium* and *Giardia*, respectively, is the presence of a unique class of microtubule associated proteins, called Microtubule Inner Proteins (MIPs) that are regularly distributed along the luminal surface of the microtubules (Nicastro *et al.*, 2006; Nicastro *et al.*, 2011; Maheshwari *et al.*, 2015; Ichikawa *et al.*, 2017; Kirima and Oiwa, 2017). The 3D structures and periodicities of MIPs have been best described in doublet microtubules (DMTs) of motile cilia where they assemble into a complex arrangement of at least 6 conserved MIP structures (Nicastro *et al.*, 2006; Nicastro *et al.*, 2011; Ichikawa *et al.*, 2017) (Fig. 1B). However, the protein composition, interactions and biological function(s) of these and other MIPs have remained elusive.

**Figure 1.**
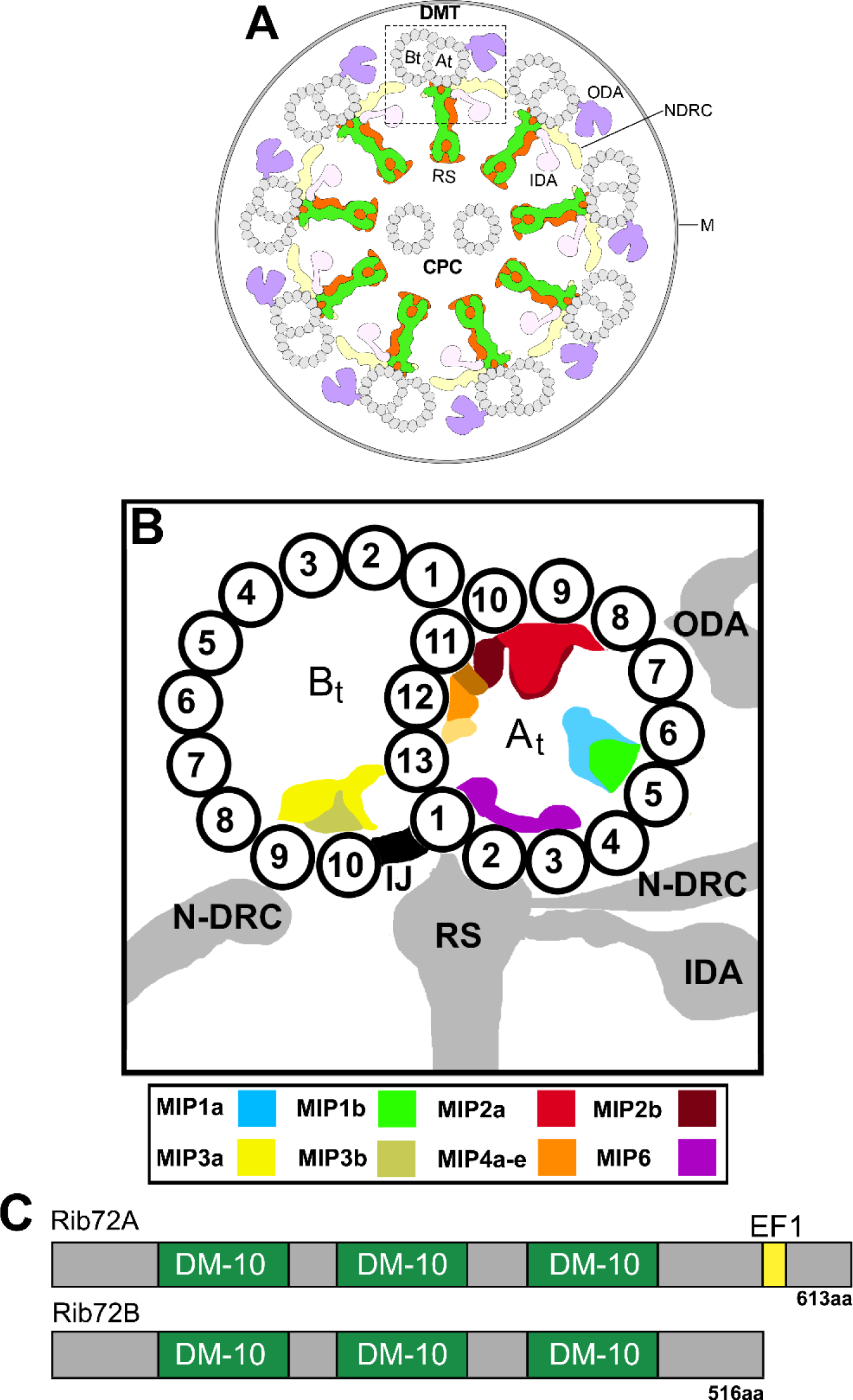
Schematic models showing the general organization of Microtubule Inner Proteins (MIPs) in the doublet microtubules, and predicted structural motifs of *Tetrahymena thermophila* RIB72A and RIB72B. (A) Simplified cross-sectional view of a cilium with nine doublet microtubules (DMTs) surrounding the central pair complex (CPC); the viewing direction is from the proximal cilium end. Outer and inner dynein arms (ODA, IDA), and the nexin-dynein regulatory complex (N-DRC) connect neighboring DMTs, whereas the radial spokes (RS) connect to the CPC. One DMT is boxed and shown larger in (B). **(B)** Schematic of a DMT with known axonemal MIPs and other microtubule-associated structures in cross-sectional view. Note that MIP5 in the B-tubule is not shown, as it is too small to be resolved by conventional cryo-ET. **(C)** Domain structures of *Tetrahymena thermophila* RIB72A and RIB72B as predicted by NCBI BLAST, showing three DM-10 domains of unknown function for both proteins, and for RIB72A a C-terminal EF-hand motif. At, A-tubule; Bt, B-tubule; IJ, inner junction; M, membrane.

Using genetic, biochemical and imaging techniques, including cryo-electron tomography (cryo-ET), we characterized two MIP candidates, RIB72A and RIB72B, in the motile cilia of *Tetrahymena thermophila.* Motile cilia typically contain a [9+2] axonemal core with 9 DMTs surrounding a central pair (CP) of singlet microtubules (Fig. 1A) (Carbajal-Gonzalez *et al.*, 2013). Each DMT consists of an A-tubule and B-tubule, with 13 and 10 protofilaments (PFs), respectively (Witman *et al.*, 1978) (Fig. 1B). Outer and inner dynein motors, and complexes that regulate the activity of the dyneins, such as the nexin-dynein regulatory complex and radial spokes, are arranged regularly along the A-tubules to generate the oscillatory movement of cilia and flagella (Gibbons, 1965; Porter and Sale, 2000; Heuser *et al.*, 2009). The 72-kDa ribbon-associated protein RIB72 was first identified as a protein associated with the “hyperstable protofilament-ribbon” a structure left after Sarkosyl extraction of sea urchin sperm flagella (Hinchcliffe and Linck, 1998). Subsequent studies showed that RIB72 is conserved from algae to humans, and tightly associates with the flagellar DMTs - possibly on the luminal side - of the unicellular algae *Chlamydomonas reinhardtii* (Ikeda *et al.*, 2003). Mutations in the human RIB72 homolog, hsEFHC1, cause juvenile myoclonic epilepsy (Suzuki *et al.*, 2004). A common feature of RIB72 homologs are three DM10 domains that are important for ciliary localization (Zhao *et al.*, 2016) and in many cases a C-terminal, Ca^2+^-binding EF-hand motif (Zhao, 2015) (Fig. 1C).

Here consistent with previous findings that localized RIB72 to *Chlamydomonas* flagella (Ikeda *et al.*, 2003), we detected two RIB72 proteins, RIB72A and RIB72B, in the basal bodies and cilia of *Tetrahymena.* Individual and double knockout mutant cells showed slower swimming speed and abnormal ciliary waveform. A null mutant-wildtype comparison using cryo-ET combined with subtomogram averaging revealed severe structural defects of the A-tubule MIP1, 4,and 6. This shows that RIB72A and RIB72B are required for the assembly of a large subset of MIP structures that may play an important role in proper ciliary motility.

## Results

### RIB72A andRIB72B both localize to basal bodies and cilia in Tetrahymena

There are two genes encoding RIB72 in the genome of *Tetrahymena thermophila: RIB72A TTHERM_00143690*) and *RIB72B (TTHERM_00584850*) (Eisen *et al.*, 2006). Previous proteomics studies revealed RIB72A protein as a basal body component in *Tetrahymena* (Kilburn *et al.*, 2007). We expressed RIB72A-mCherry fusion protein in *Tetrahymena* cells co-expressing either the basal body marker centrin (GFP-CEN1) or α-tubulin (GFP-ATU1). Fluorescence microscopy revealed that RIB72A-mCherry localized to basal bodies (Fig. 2A-C) and along the entire ciliary length (Fig. 2D-F).

**Figure 2.**
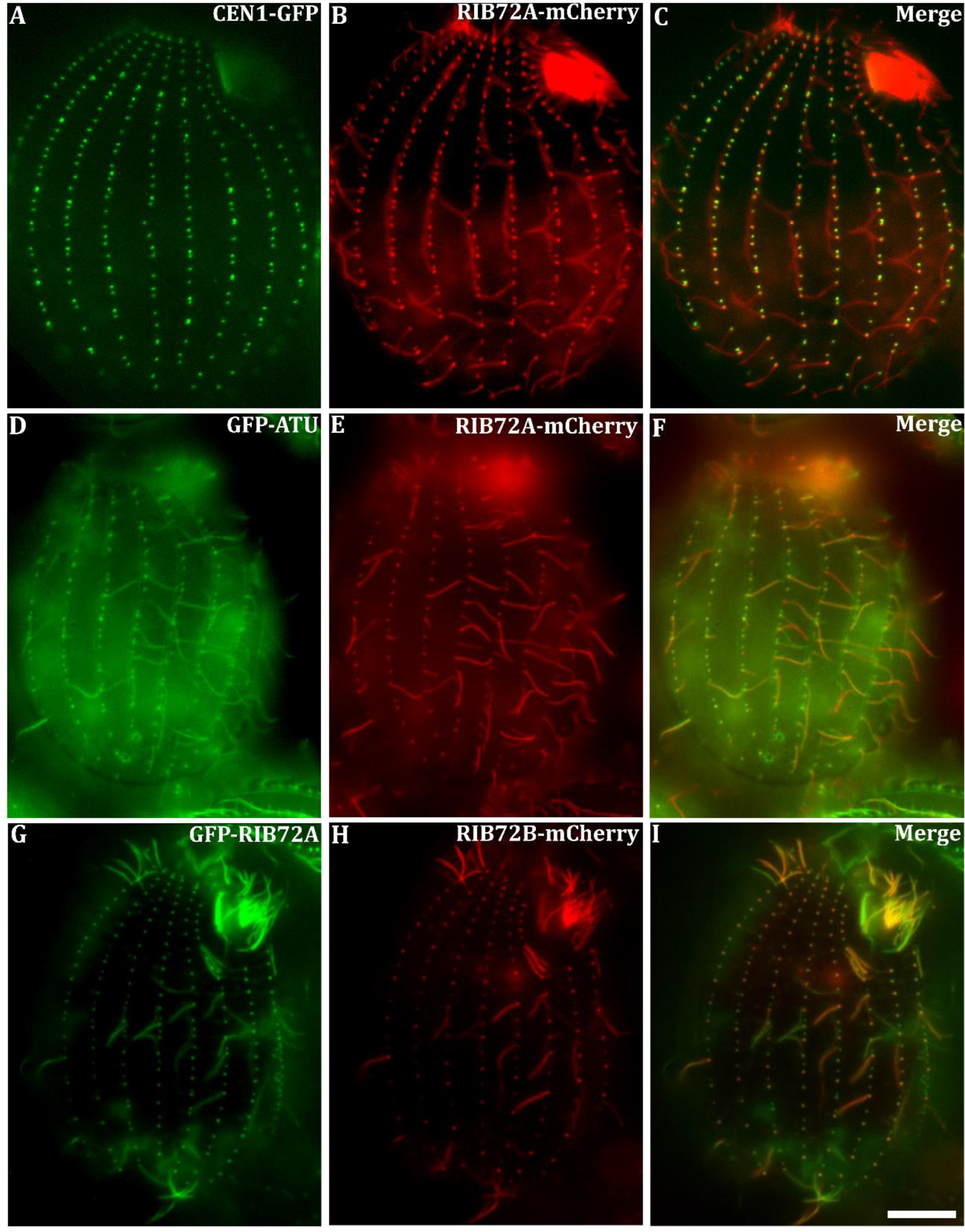
Fluorescence images reveal that *Tetrahymena* RIB72A and RIB72B co-localize to cilia and basal bodies. Fluorescence light microscopy images of fixed *Tetrahymena* cells expressing RIB72A or RIB72B tagged by fluorescent proteins along with basal body and microtubule markers. **(A-C)** RIB72A-mCherry (B) coexpressed with the basal body marker GFP-CEN1 (A) shows colocalization at basal bodies (C) in a WT background. **(D-F)** RIB72A-mCherry (E) coexpressed with the microtubule marker GFP-ATU1 (D) showing colocalization along the ciliary length (F) in a WT background. **(G-I)** Coexpressed GFP-RIB72A (G) and RIB72B-mCherry (H) showing colocalization at basal bodies and cilia (I) in a WT background. Cilia displaying signal from a single tagged Rib72 protein moved during imaging as live cells were used to limit fixation artifacts. Scale bar: 10 μm.

We constructed *Tetrahymena* knockout (KO) strains lacking either *RIB72A, RIB72B*, or both genes. GFP-RIB72A expressed in the *RIB72B-KO* background localized to both cilia and basal bodies (Fig. S1A), suggesting that RIB72A assembles in these organelles independently of RIB72B. Also, RIB72B-mCherry localized to both cilia and basal bodies in the absence of RIB72A (Fig. S1 A, B). Co-expression of GFP-RIB72A and RIB72B-mCherry showed co-localization of the two proteins in most but not all cilia suggesting that RIB72A and RIB72B co-exist in at least a subset of cilia (Fig. 2G-I).

### Loss of RIB72A and/or RIB72B decreases swimming speed and phagocytosis rate in Tetrahymena

The swimming speeds of *RIB72A-KO* (~160 μm/sec) and *RIB72B-KO* (~106 μm/sec) cells were reduced to 2/3 and half of the WT velocity (245 μm/sec), respectively (Fig. 3A). The swimming velocity of the double-knockout *RIB72A/B-KO* (~112 μm/sec) was about as low as that of *RIB72B-KO* (Fig. 3A).

**Figure 3.**
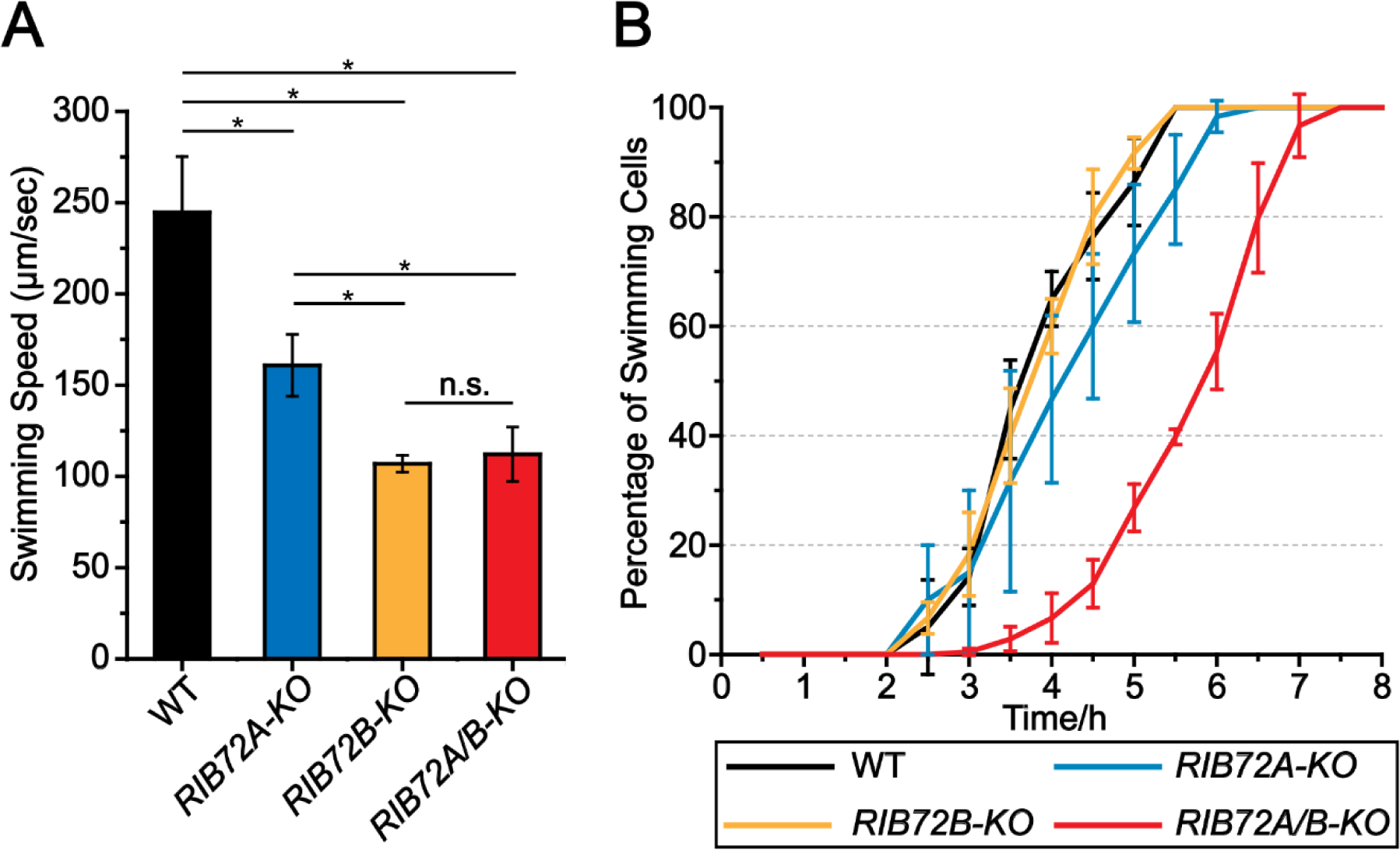
Gene knockouts of *RIB72A* and *RIB72B* decrease swimming velocity of *Tetrahymena* cells. (A) Loss of RIB72A or RIB72B decreases swimming velocity as compared to WT (100%): *RIB72A-KO* 66%, *RIB72BKO* 44%, *RIB72A/B-KO* 46% (n=20 cells per stain). Error bars represent standard deviations and asterisks indicate statistical significance at p<0.01 (one way ANOVA). n.s. stands for non-significant, p>0.05. **(B)** The percentage of cells recovering motility (>25 μm/sec) after deciliation was recorded.

*Tetrahymena* use oral cilia to move particles into the oral groove for feeding through phagocytosis. We conducted a phagocytosis assay by adding India ink as “food” to the growth media and scoring the number of ink-containing food vacuoles per cell. The food vacuoles were smaller and less numerous in both *RIB72B-KO* and *RIB72A/B-KO* (Fig. S1 C-F). These results show that RIB72A and especially RIB72B are important for the functionality of oral cilia.

Following deciliation by a pH shock, all strains, WT and the three KO mutants, were able to regenerate cilia as evidenced by the recovery of cell motility, but *RIB72A-KO* and especially the double mutant *RIB72A/B-KO* restored motility at a slower rate than WT (Fig. 3B), indicating that *RIB72A-KO* and double knockout cells either assemble slower or fewer cilia following deciliation.

### High speed video imaging reveals abnormal waveform and decreased beating frequency in the three RIB72A/B knockout strains

To analyze cilia beating in live cells, we collected high-speed videos of forward-swimming cells that were partially restrained inside microfluidic channels (Videos S1-S5). Using an image processing workflow that was originally developed for *Paramecium* (Funfak *et al.*, 2015), we extracted video segments that contain the zone of beating cilia around the entire cell circumference as outlined in Fig. 4A. These videos were used to produce chronographs along a single line positioned slightly above the cell body surface (Fig. 4B-E, also see Fig. S2 for technical details). In the chronograph of a wild-type cell, short greyscale diagonal lines represent traces of individual cilia as they undergo their power stroke (Fig. 4B). In the recovery stroke, the cilia are too close to the cell surface to be visible (Fig. S2). The chronograph shows the time (y-axis) that it takes for a cilium occupying a particular position on the cell surface (x-axis) to pass through the field of view (examples are shown by the red arrows in Fig. 4B-E). Wild-type cilia were uniformly active along the entire cell surface and were beating with an average frequency of 38 Hz (+/− 3.2, n=45 cilia, 9 cells) (Fig. 4B, F). When a cilium stalls, this produces a vertical or wavy line on the chronograph (Fig S2). The *RIB72A-KO* chronogram contained a few vertical or wavy lines consistent with stalled or pivoting cilia (red box in Fig. 4C), while such cilia were not apparent in the *RIB72B-KO* chronograph (Fig. 4D). The stalled cilia were very prominent in the double knockout chronograph (red boxes in Fig. 4E). In the single and double knockouts, the cilia that were not stalled showed reduced beat frequencies. The *RIB72A-KO* and *RIB72B-KO* cilia beat at 29 and 28 Hz, respectively (23% and 27% reduction compared to wild type) and the *RIB72A/B-KO* cilia beat at 23 Hz (39% reduction). The inverse slope of each diagonal line in the chronograph represents the speed of power stroke. The power stroke in the double knockout cilia was 43% slower as compared to the wild type (Fig. S3A). Thus, the reduced beat frequency in *RIB72A/B-KO* is at least partly due to the slower movement of the cilium during the power stroke.

**Figure 4.**
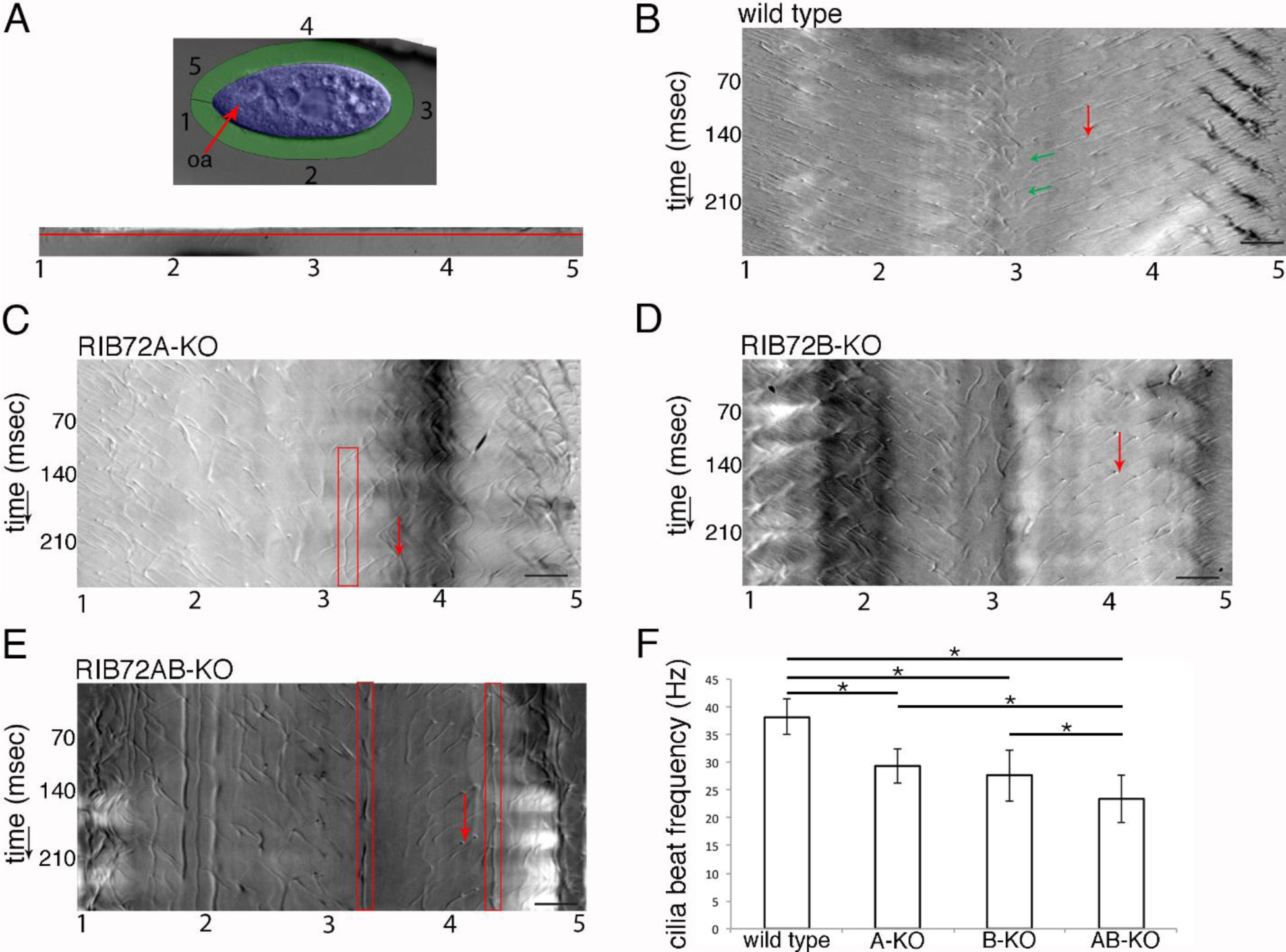
Chronographs based on high-speed video imaging show that loss of either RIB72A or RIB72B or both proteins strongly affects the motility of cilia. (A) Outline of the image analysis workflow, based on Funfak et al. (2014) as applied here to *Tetrahymena* (see Fig. S2 for details). As shown in the top panel, the first frame of a video was used to mark the cell body (blue) and cilia (green) regions. Numbers correspond to the positions along the cell circumference starting at the cell’s anterior tip and going around the cell in a counterclockwise direction. The bottom panel shows a single frame of a movie of an extracted and unwrapped cilia region. Numbers 1–5 correspond to the positions marked around the cell circumference as shown in the top panel. The red line shows the position which was plotted over time to produce the chronograph seen in (B). **(B-E)** Chronographs of the ciliary zones from wild-type (B) (video S1), single (C, D) (video S2-S3) and double knockout cells (E) (video S4). The numbers on the x-axis correspond to the positions around the cell circumference as indicated in panel (A), the y-axis represents time. Red arrows show examples of single beat duration measurements as distances between two consecutive diagonal gray-scale lines representing the same cilium in consecutive beat cycles. Green arrows show the power stroke of a single cilium. Note the presence of stalled or pivoting cilia indicated by red boxes in C and E. **(F)** Average beat frequencies in the wild-type (38.1 + 3.2 Hz, n cilia = 45, n cells = 9), *RIB72A-KO* (29.3 + 3.1 Hz, n cilia = 48, n cells = 8), *RIB72B-KO* (27.6 + 4.6 Hz, n cilia = 64, n cells = 9) and *RIB72AB-KO* cells (23.3 + 4.3 Hz, n cilia = 50, n cells = 8) cells. Error bars represent standard deviations and asterisks indicate statistical significance at p<0.001 (one way ANOVA). Abbreviations: oa, oral apparatus. Scale bars are 10 microns.

The metachronal waves were visible near the anterior end of the wild type chronograph, as dark diagonal lines (Fig. 4B) (Funfak *et al.*, 2015). The metachronal waves were visible but less frequent in the single KO strains but completely absent in the double knockout cells (Fig. 4B-E). Thus, the loss of RIB72A/B also affects the coordination of beating between cilia. An analysis of the successive frames of extracted cilia videos for the double knockout cells revealed abnormalities in the shape of individual cilia. Specifically, during the power stroke, wild-type cilia remained relatively straight, while some of the *RIB72A/B-KO* cilia appear excessively curved (Fig. S3B). Interestingly, the curving occurred halfway into the power stroke and in the opposite direction to the bend that normally forms during the recovery stroke. Furthermore, some *RIB72A/B-KO* cilia had a kink near the distal tip (Video S5). We conclude that the loss of RIB72A and RIB72B reduces the beat frequency, increases the incidence of cilium stalling, reduces the metachronal activity and alters the shape of the cilium during the power stroke.

### Cryo-ET reveals defects in the MIP1, MIP4 and MIP6 structures in *RIB72A-KO* and *RIB72B-KO*, and additive defects in the double knockout

We performed cryo-ET and subtomogram averaging of the 96-nm axonemal repeats and compared the 3D structures between WT, *RIB72A-KO, RIB72B-KO* and *RIB72A/B-KO* to determine the effects of RIB72A and/or RIB72B removal on the ciliary structure. The axonemal microtubules themselves (DMTs and CPC) and all axonemal structures that bind to the exterior of the microtubules, such as the dyneins and regulatory complexes, appeared fully assembled in all four strains (Fig. S4). In contrast, the mutant averages revealed clear defects in several MIP structures on the luminal side of the A-tubule (Figs. 5-7). Previous cryo-ET studies named the major MIP densities in ciliary DMTs as MIP1-6 based on their association with certain subsets of DMT PFs (Fig. 1B). Absence of RIB72A and RIB72B led to defects in different subcomplexes of MIP structures, namely of MIP1 (PF A5-A7) (Fig. 5), MIP4 (PF A10-A13) (Fig. 6) and MIP6 (PF A1-4) (Fig. 7) (summarized in Table 1). In the double mutant *RIB72A/B-KO* these defects were additive and thus most severe (Figs. 5-7) (summarized in Table 1).

**Figure 5.**
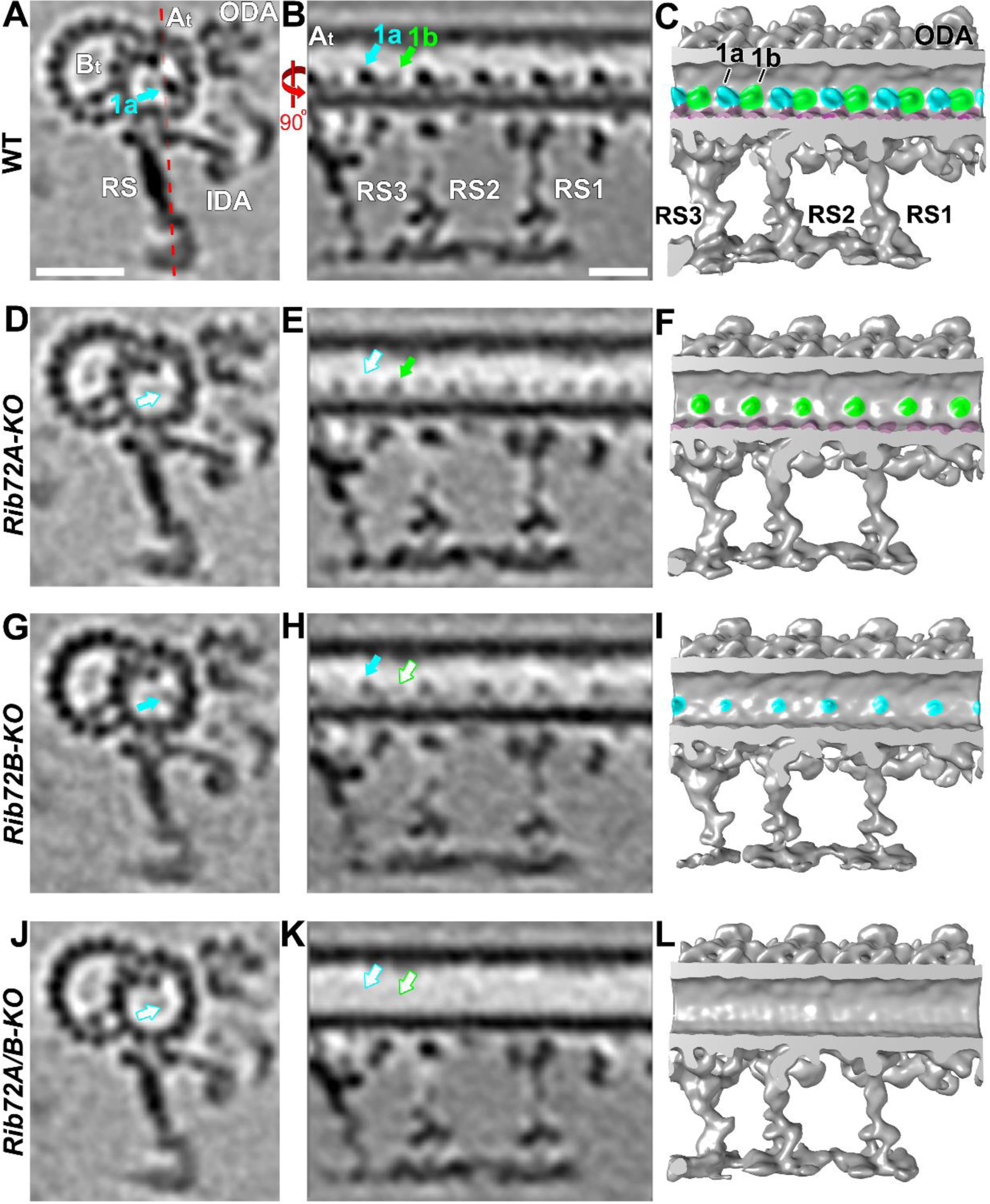
Losses of RIB72A and RIB72B affect the MIP1 structure differently. (A-L) Averaged tomographic slices (A, B, D, E, G, H, J, K) and isosurface renderings (C, F, I, L) in cross-sectional (A, D, G; from proximal) and longitudinal view (B, C, E, F, H, I; proximal on the right) of *Tetrahymena* axonemal repeat units of *RIB72A-KO* (D-F), *RIB72B-KO* (G-I), and the double mutant (J-L) showed different defects in the MIP1a (blue) and/or MIP1b (green) structure compared to WT (A-C). The *RIB72A-KO* mutant is missing MIP1a (open blue arrow), but not MIP1b (solid green arrow); the *RIB72B-KO* mutant lacks MIP1b (open green arrow), but not MIP1a (solid blue arrow). The double KO shows both defects, i.e. MIP1a and MIP1b are missing (open blue and green arrows). Red dotted line in (A) indicates slice position of longitudinal tomographic slices, e.g. in (B). Other labels: At, A-tubule; Bt, B-tubule; IDA, inner dynein arms; ODA, outer dynein arms; RS, radial spokes. Scale bars: 25 nm (A, valid also for D, G), 16 nm (B, valid also for E, H).

**Table 1.** Summary of MIPs present or absent in WT and here studied mutants

| MIP components<br>(~estimated MW) | Strain |  |  |  |
| --- | --- | --- | --- | --- |
|  | WT | *RIB72A-KO* | *RIB72B-KO* | *RIB72A/B-KO* |
| 1a (~55) | + | - | + | - |
| 1b (~20) | + | + | - | - |
| 4b (~20) | + | - | + | - |
| 4d (~20) | + | - | + | - |
| 4e (~25) | + | +/- | - | - |
| 6a (~25) | + | + | - | - |
| 6b (~18) | + | - | + | - |
| 6c (~15) | + | - | + | - |
| 6d (~20) | + | + | - | - |

+ indicates that the component is present; +/- indicates that the component reduced; - indicates that the component is lacking. Molecular mass is indicated in parentheses (in kDa).

MIP1 consists of a larger MIP1a and an intervening shorter MIP1b subunit, and together the MIP1a/b minimum repeat has a 16-nm periodicity (Fig. 5A-C). The density corresponding to MIP1a was missing in *RIB72A-KO*, whereas *RIB72B-KO* lacked MIP1b (Fig. 5D-I), and the double mutant *RIB72A/B-KO* lost the entire density of MIP1 (Fig. 5J-L). From the subtomogram averages, we estimated the molecular mass of the subunit MIP1a to be ~50 kDa and that of MIP1b ~20 kDa.

MIP4 consists of multiple densities that are attached to the A-tubule-luminal side of the DMT midpartition (PFs A10-A13), which was also shown to represent most of the hyperstable PF-ribbon (Nicastro *et al.*, 2011; Linck *et al.*, 2014). Here, we could distinguish five sub-densities of the *Tetrahymena* MIP4, namely MIP4a-e, which showed an overall minimum repeat periodicity of 48 nm along PF A12 (MIP4a-d) and PF A11-12 (MIP4e) (Fig. 6C). Based on the tomographic data the sizes of these MIP4 sub-densities were estimated to be ~20 kDa each (Fig. 6A-C). The structural defects of MIP4 were relatively mild in *RIB72B-KO* axonemes that lacked only MIP4e (Fig. 6G-I). In contrast, the defects were similarly strong in *RIB72A-KO* and *RIB72A/B-KO* that were missing MIP4b and 4d, and were reduced in (Fig. 6D-F) or missing MIP4e (Fig. 6J-L), respectively. The densities of the A-tubule MIP2 along PF A8-A12 remained completely unchanged in the mutants (Fig. 6).

**Figure 6.**
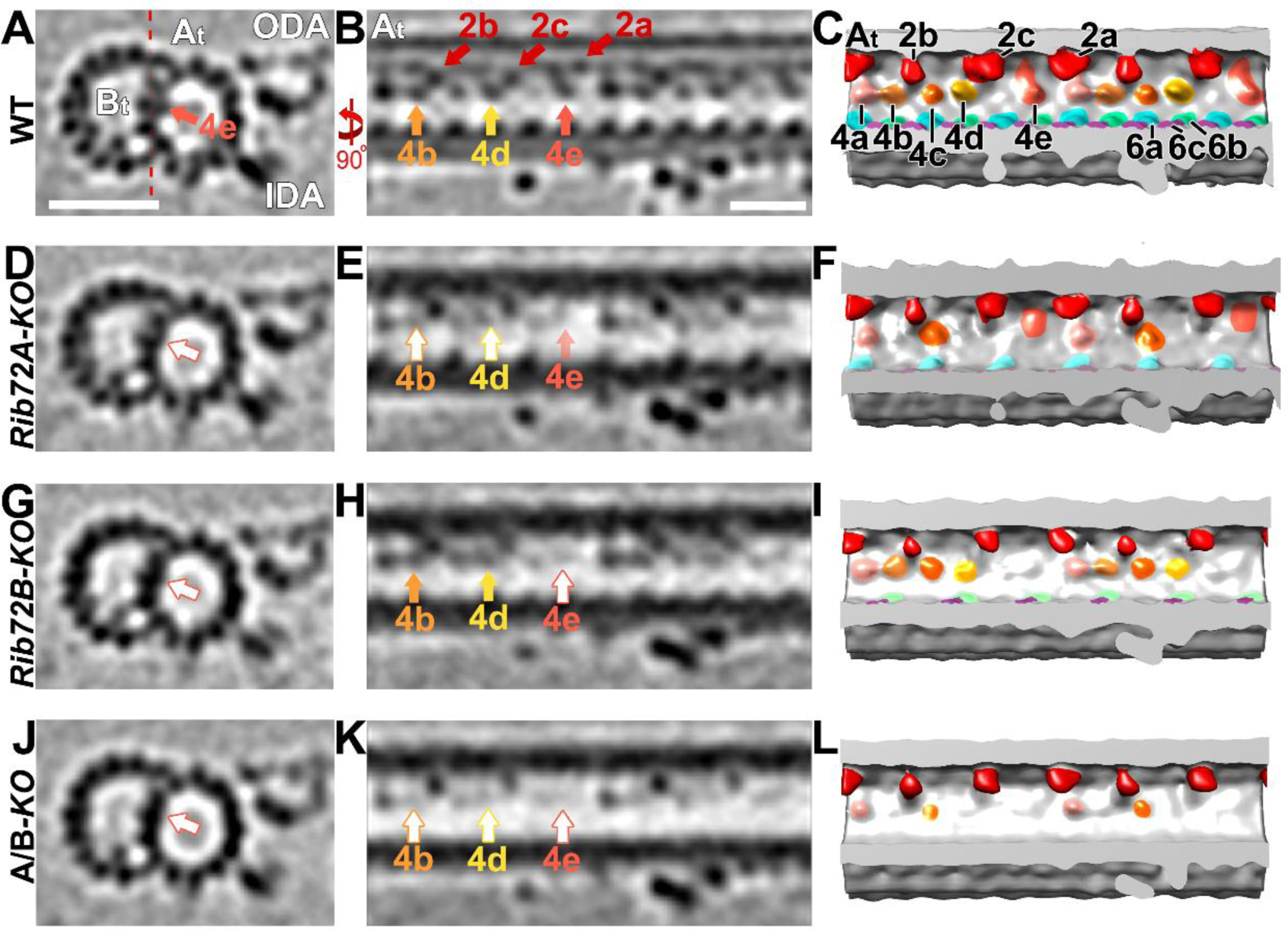
Loss of RIB72A/B affects multiple MIP4 subunits. (A-L) Tomographic slices (A, B, D, E, G, H, J, K) and isosurface renderings (C, F, I, L) in cross-sectional (A, D, G, J; from proximal) and longitudinal view (B, C, E, F, H, I, K, L; proximal on the left) of *Tetrahymena* axonemal repeat units. *RIB72A-KO* (D-F), *RIB72B-KO* (G-I), and the double mutant (J-L) have various defects in the MIP4 structures (4a, rose; 4b, light orange; 4c, dark orange; 4d, yellow; 4e tomato) as compared to WT (A-C); missing/reduced structures are indicated by open arrows outlined by the respective colors for MIP4 subunits. The *RIB72A* mutant has defects in the MIP4b (light orange arrow), MIP4d (yellow arrow), and a slight defect in MIP4e (tomato arrow). The *RIB72B* mutant has defects only in the MIP4e subunit, and no others. The defects in the double KO were additive, i.e. MIP4b, 4d, and 4e are missing. Red dotted line in (A) indicates slice position of longitudinal tomographic slices, e.g. in (B). Other labels: At, A-tubule; Bt, B-tubule; IDA, inner dynein arms; ODA, outer dynein arms; RS, radial spoke; red, MIP2a/b. Scale bars: 25 nm (A, valid also for D, G, J), 16 nm (B, valid also for E, H, K).

MIP6 was not originally described, but visible in higher resolution single-particle cryo-EM reconstruction spanning PFs A1-4 with 8 nm periodicity (Maheshwari *et al.*, 2015). In our studies, by comparing the structural defects in *RIB72A-KO, RIB72B-KO* and *RIB72A/B-KO* to wildtype axonemes, we found that MIP6 was composed of at least 4 different subunits, namely MIP6a-d, that ranged in size from ~15-25 kDa each (Fig. 7). MIP6a and 6b alternated along PF A1 with an overall periodicity of 16 nm, whereas the MIP6c and 6d densities formed a continuous ridge that spanned PFs A2-3 with an overall periodicity of 8 nm (Fig. 7A-C). The structural defects in the single KO mutants were complementary, i.e. *RIB72A-KO* was missing MIP6b and MIP6c (Fig. 7D-F), whereas *RIB72B-KO* lacked MIP6a and MIP6d (Fig. 7G-I). Similar to MIP1, the defects in the double mutant *RIB72A/B-KO* were additive, i.e. the entire MIP6 structure was absent (Fig. 7 J-L).

**Figure 7.**
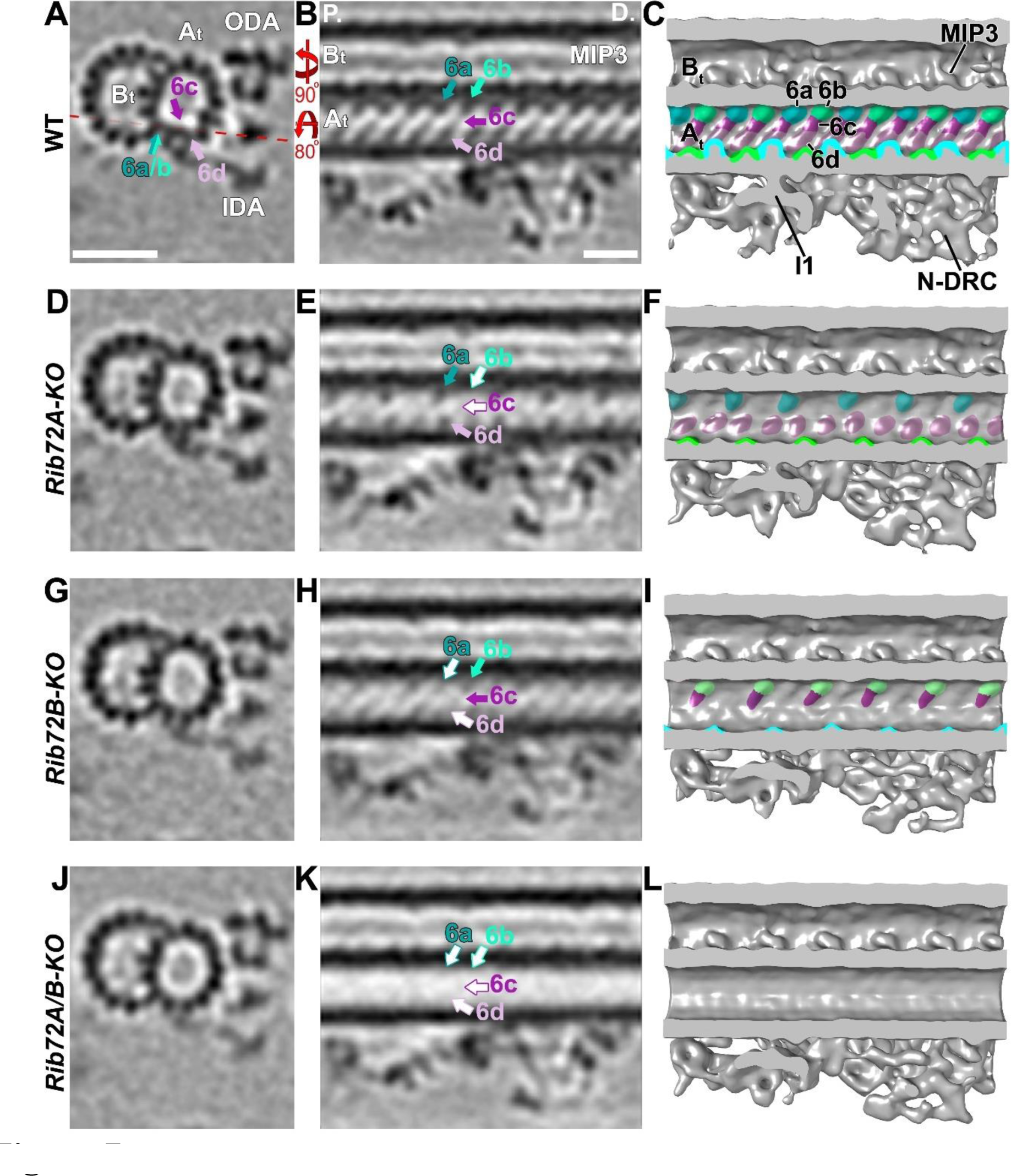
Loss of RIB72A/B affects the MIP6 structure and shows that MIP6 is composed of multiple subunits. (A-L) Tomographic slices (A, B, D, E, G, H, J, K) and isosurface renderings (C, F, I, L) in cross-sectional (A, D, G, J; from proximal) and longitudinal bottom view (B, C, E, F, H, I, K, L; proximal on the left) of *Tetrahymena* axonemal repeat units. *RIB72A-KO* (D-F), *RIB72B-KO* (G-I), and the double mutant (J-L) each have defects in the MIP6 structures (6a, blue; 6b, aqua; 6c, purple; 6d, rose) as compared to WT (A-C); missing structures are indicated by open arrows outlined by the respective colors for MIP6 subunits. The *RIB72A* mutant lacked MIP6b (aqua arrow) and MIP6c (purple arrow), whereas the *RIB72B* mutant was missing only MIP6a (blue arrow) and MIP6d (rose arrow). The defects in the double KO were additive, i.e. all four MIP6 subunits were missing. Red dotted line in (A) indicates slice position of longitudinal tomographic slices, e.g. in (B). Other labels: At, A-tubule; Bt, B-tubule; I1, I1 dynein; IDA, inner dynein arms; MIP3; N-DRC, nexin dynein regulatory complex; ODA, outer dynein arms. Scale bars: 25 nm (A, valid also for D, G, J), 16 nm (B, valid also for E, H, K).

### RIB72B-GFP rescue shows additional density in the area of MIP4

To test if RIB72B protein was a structural component of the RIB72B-dependent MIP-complex or only required for its assembly, we imaged axonemes from a *RIB72B-KO* rescue-strain that expressed N-terminally tagged RIB72B-GFP. The swimming speed of the RIB72B-GFP rescue strain was similar to that of WT (Fig. S1H), and all structural MIP defects of *RIB72B-KO* (MIPs 1b, 4e, 6a, 6d) were rescued to resemble the WT structure (Fig. 8). Moreover, the resolution of the subtomogram averages was sufficiently high to visualize the additional density corresponding to GFP (27 kDa) at the inner surface of the A-tubule PF A12 and close to the proximal end of MIP4e, demonstrating that RTB72B is a structural component of the RTB72B-dependent MIP-complex with its C-terminal domain likely located at MIP4e (Fig. 8F, L). We also attempted to visualize GFP-tagged RIB72A, however, in the studied strains neither full structural rescue of the RIB72AKO nor GFP-tagged RIB72A was observed in the axonemal averages.

**Figure 8.**
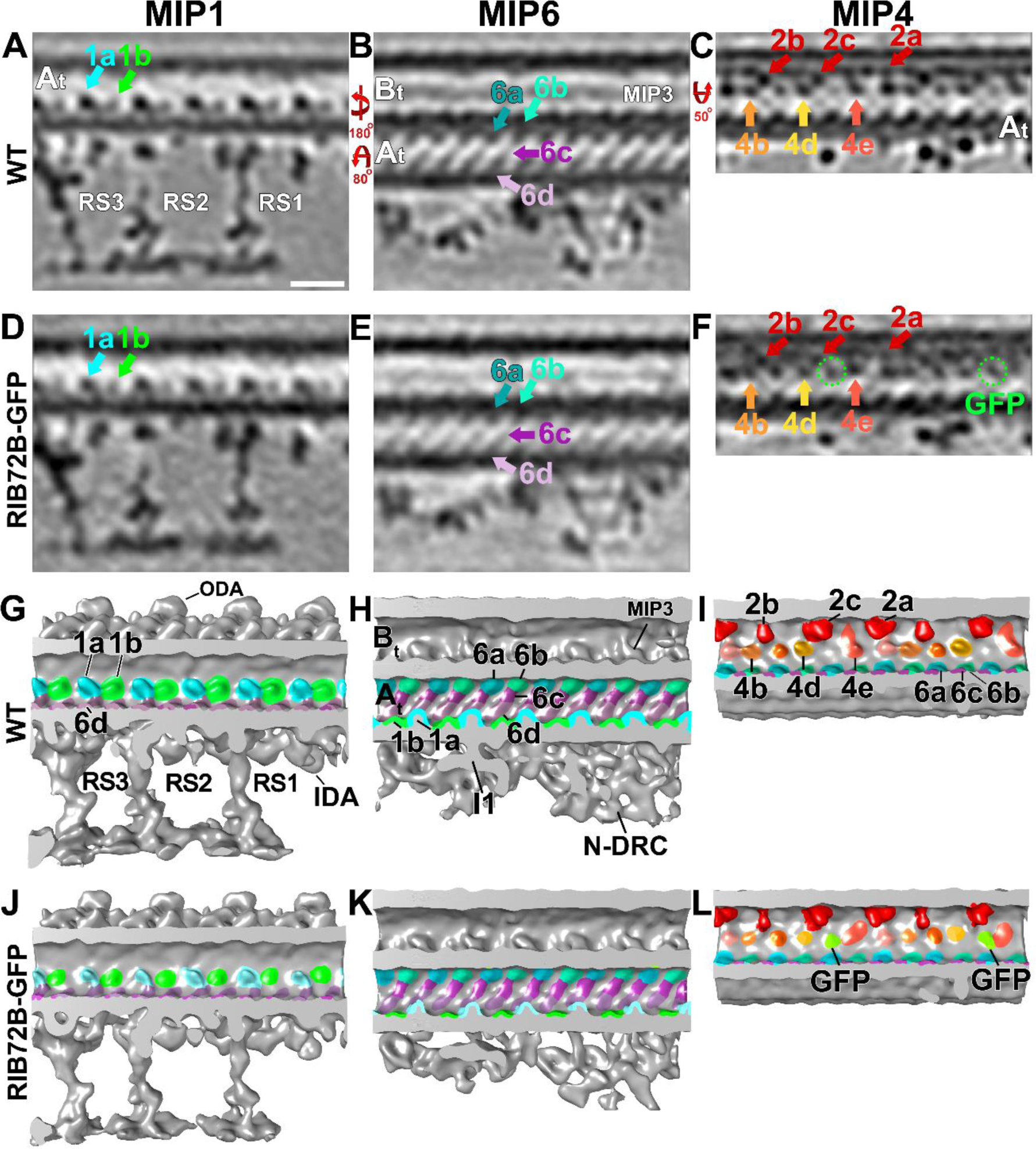
*RIB72BKO* axonemes rescued with RIB72B-GFP show no structural defects and an additional tag density. (A-L) Tomographic slices (A-F) and isosurface renderings (G-L) in longitudinal views (in A, D, G, J proximal is on the right; in all remaining panels proximal is on the left) of *Tetrahymena* axonemal repeat units. The axonemal average from *RIB72B-KO* rescued with RIB72B-GFP (D-F, J-L) showed the same structures as seen in WT (A-C, G-I), i.e. the mutant defects were rescued: MIP1b (green arrow and structures in D, J), MIP6a and 6d (dark blue and rose arrow and structures in E, K), and MIP4e (tomato arrow and structures in F, L). In addition, the extra density of the GFP-tag was visible in the vicinity of MIP4e in the rescued strain (green circles and densities in F, L). Slice positions of the tomographic slices are the same as in Figs. 5 (for MIP1), 6 (for MIP6), and 7 (for MIP4); MIP structures are colored as described in Fig. 5–7. Other labels: At, A-tubule; Bt, B-tubule; I1, I1 dynein; IDA, inner dynein arms; MIP3; N-DRC, nexin dynein regulatory complex; ODA, outer dynein arms; RS, radial spokes; GFP, green fluorescent protein. Scale bar: 16 nm (valid A-F).

## Discussion

### RIB72A and RIB72B are conserved MIPs required for the assembly of two independent MIP supramolecular complexes inside the ciliary A-tubule

Our subtomogram averages clearly showed that several A-tubule MIP structures were missing or reduced in the absence of RIB72A and/or RIB72B. In addition, our localization of the additional GFP-tag density at PF A12 in the RIB72B-GFP rescue strain demonstrated that RIB72B itself is a component of the RIB72B-dependent MIP-complex. This is consistent with previous studies, such as i) identification of RIB72 as a component of the hyperstable ribbon extracted from Sarkosyl-treated axonemes (Patel-King *et al.*, 2002; Linck *et al.*, 2014), ii) negative-staining immuno-EM with anti-RIB72 antibodies that labeled partially splayed or degraded A-tubules more frequently than intact A-tubules (Ikeda *et al.*, 2003), and iii) a more recent cryo-immunogold-ET study with anti-SpRIB74 (the RIB72 homolog in the sea urchin *Strongylocentrotus purpuratus)* that detected gold labels along the side of the hyperstable ribbon facing the A-tubule lumen, again suggesting that spRIB74 localizes to the luminal surface of PF A11-12-13-1 (Linck *et al.*, 2014). However, previous studies were hampered by limited efficiency of antibody labeling and reduced resolution of negative staining immuno-EM, the long labels with primary and secondary antibodies (20–25 nm length), and by the reduced structure preservation due to the need for partial disassembly of the DMTs to allow antibody-access to the luminal microtubule surface. Here we provided direct visual evidence that RIB72B of *Tetrahymena* is a MIP bound to the luminal surface of the A-tubule, and that absence of RIB72A and/or RIB72B causes severe structural MIP defects. These data suggest that RIB72A and RIB72B are crucial for the stable assembly of two independent supramolecular MIP-complexes that together cover the luminal surface of about half of the A-tubule diameter. The RIB72A-dependent and RIB72B-dependent MIP-supra-complexes were complementary and composed of [MIPs 1a, 4b, 4d, (4e), 6b, 6c] and [MIPs 1b, 4e, 6a, 6d], respectively (Table 1; Fig. 9).

**Figure 9.**
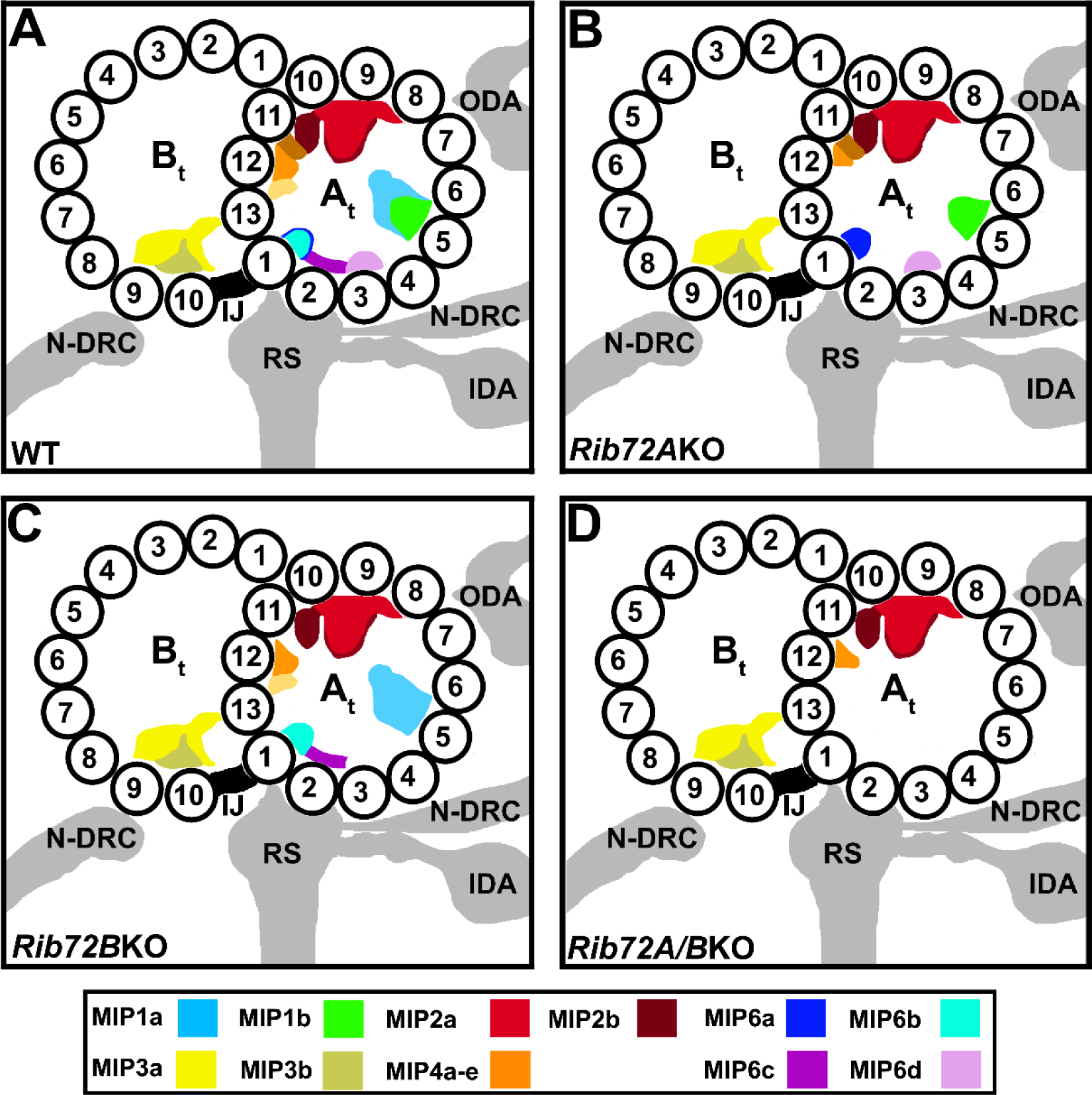
Summary schematics of MIP locations in the DMTs of WT (A), RIB72A-KO (B), *RIB72B-KO* (C) and *RIB72A/B-KO* (D) cilia. The MIP coloring as shown in the color legend. Other labels: At, A-tubule; Bt, B-tubule; IDA, inner dynein arms; IJ, inner junction; N-DRC, nexin dynein regulatory complex; ODA, outer dynein arms; RS, radial spokes; 1–13, PF numbers.

The observed structural defects in each single and the double KO mutant seemed to be larger than the molecular size of a single copy of the 72-kDa RIB72A and/or 61-kDa RIB72B proteins. Based on the missing densities in our subtomogram averages of *RIB72A/B-KO*, we estimated the total mass of the RIB72A-dependent and RIB72B-dependent MIP-supra-complexes combined to be ~240 kDa. This is nearly twice of the sum of RIB72A and RIB72B (133 kDa), suggesting that either multiple copies of each RIB72 proteins are involved, or that additional, unidentified proteins contribute to these MIP-supra-complexes. A recent biochemical and immuno-gold labeling study suggested that the EF-hand containing protein FAP85 is an A-tubule MIP and may be associated with the MIP1 structure in *Chlamydomonas* flagella (Kirima and Oiwa, 2017).

### RIB72A and RIB72B likely form two independent MIP supramolecular complexes inside the A-tubule of basal bodies

Using cryo-ET we showed that both RIB72A and RIB72B are axonemal MIPs. However, our fluorescence light microscopy localized both proteins not only to the ciliary length, but also to the basal bodies. Previous fluorescence light microscopy of labeled RIB72 homologs in other species, such as *Chlamydomonas*, also demonstrated RIB72 localization along the length of axonemes, but not to basal bodies (Ikeda *et al.*, 2003). This suggests that both proteins are also MIPs in the basal bodies, which is consistent with a previous study that identified the *Tetrahymena* proteins, RIB72A/Bbc73 and RIB72B/Bbc60, in the basal body proteome (Kilburn *et al.*, 2007). Although previous cryo-ET studies of basal bodies and centrioles were limited in their resolution, because of technical challenges due to their shorter length and accessibility/need for isolation from the cell body, some MIP densities were also observed in the triplet microtubules of these organelles (Guichard *et al.*, 2012; Li *et al.*, 2012). Together this suggests that at least some, if not all, MIP structures described for ciliary DMTs, including the highly conserved RIB72A-dependent and RIB72B-dependent MIP-supra-complexes in the A-tubule, are also assembled in the A- and B-tubules of basal bodies.

### Possible functional role(s) of ciliary MIPs

The overall structures and periodicities of MIPs inside DMTs appears conserved from algae to human (Nicastro *et al.*, 2011; Lin *et al.*, 2014), suggesting that the biological function(s) of MIPs in motile cilia is conserved. Previous studies have hypothesized various functional roles for ciliary MIPs, such as stabilization of DMT, anchoring of axonemal complexes to the outer surface, and regulation of beat frequency and/or ciliary waveform (Nicastro *et al.*, 2011; Linck *et al.*, 2014; Ichikawa *et al.*, 2017), but so far little experimental evidence exists to test these hypotheses. Here we have studied ciliary functions by phenotypically comparing wildtype and *Rib72A/B-KO* mutants, including swimming velocity, phagocytosis efficiency, and ciliary waveform and frequency. Our results showed that the MIP defects – even though all other structures seemed to have fully assembled in the axoneme – caused abnormal ciliary motility. Specifically, high-speed video imaging of beating cilia revealed decreased beating frequency and unusual waveform with occasional distal cilia kinking, demonstrating that RIB72A and RIB72B are important for the proper ciliary beating.

MIPs are a common feature of highly stable microtubules that experience mechanical stress. In contrast to regular cytoplasmic microtubules, DMTs are highly stable and cold-resistant (Andrieux *et al.*, 2002). In motile cilia, the DMTs provide a stable scaffold for many associated complexes while being exposed to vigorous bending with ~40Hz frequency, i.e. the bending direction changes every ~8 msec. In the malaria-causing pathogen *Plasmodium*, the MIPs are proposed to stabilize the subpellicular microtubules to provide the sporozoites with a highly elastic, yet stable, cytoskeleton that remains intact while the cells undergo extreme bending as they glide and squeeze through tissue barriers (Cyrklaff *et al.*, 2007). Attachment of the diarrhea-causing parasite *Giardia lamblia* to the host’s intestinal epithelium is mediated by a suction-type mechanism caused by the ventral disc, a large spiral-shaped array of stable and MIP-containing microtubules (Schwartz *et al.*, 2012). The microtubule-stabilizing drug taxol and its derivatives also bind to the inside of the microtubule wall (Nogales *et al.*, 1999). Thus, the inner scaffold formed by MIPs could add physical interactions that strengthen tubulin dimer and protofilament coherence, thereby increasing microtubule stability and maybe regulating microtubule elasticity.

However, despite severe MIP defects, especially in the double *RIB72A/B-KO* mutant, we did not observe large scale DMT defects by cryo-ET in inactive (ATP-free) isolated axonemes, such as partially depolymerized microtubules or holes in PFs, neither in the raw tomograms nor the subtomogram averages. It is possible that smaller defects in the microtubule lattice were present, but averaged out in the subtomogram averages. However, imaging of actively beating mutant cilia revealed abnormally high-curved cilia during the power stroke and in the double KO there were occasional kinks (Fig. S3B) that could be a sign of a broken axoneme. Smaller axonemal pieces that were broken off would likely get lost during axoneme isolation. The abnormal curvature and kinks in mutant cilia suggests that RIB72A and RIB72B probably contribute to the stability and possibly the elasticity of the axoneme in actively beating cilia.

Another possible function of MIPs could be regulation during cilia movement. One proposed regulatory mode is the interaction of MIPs with axonemal complexes on the outside of DMTs, contributing to their regulation. In fact, protease digestion experiments in which a ~10- kDa fragment was cleaved from *Chlamydomonas* RIB72 suggested that this fragment was likely to reside on the outer microtubule surface and was therefore accessible to protease (Ikeda *et al.*, 2003; Ichikawa *et al.*, 2017). This fragment could interact with associated axonemal complexes, such as the regulatory I1 dynein or the calmodulin and spoke associated complex (CSC), and through these interactions regulate cilia motility. Importantly, a recent high resolution study showed MIPs projecting through the DMT and interacting with proteins on the exterior of the DMT (Ichikawa *et al.*, 2017). Interestingly, the EF-hand calcium-binding domain at the C-terminus of RIB72A is highly conserved among various species (Ikeda *et al.*, 2003). In *Chlamydomonas*, the flagella show a Ca^2+^-dependent waveform conversion, and the sensitivity of RIB72 to trypsin digestion depends on the Ca^2+^ concentration (Ikeda *et al.*, 2003). Axoneme curvature has also been proposed as a signal that switches dynein activity on/off on opposite sides of the flagellum to generate the undulating motion typical of cilia (Satir and Matsuoka, 1989). If the MIPs change the stability/elasticity properties of the DMTs, this could also result in delayed or lack of proper signaling to coordinate dynein activities in beating cilia. Combined with our highspeed imaging results showing abnormal beating waveform and frequencies of MIP-depleted cilia, including stalling and pivoting, RIB72A and RIB72B might also be important for signaling and regulation of ciliary motility.

We demonstrated that RIB72A and RIB72B are ciliary MIPs that are required for proper ciliary motility. These proteins may contribute to DMT stability and elasticity, and possibly to the regulation of ciliary beating.

## Materials and Methods

### Construction of gene knockout strains

Native expression of C-terminal fusion proteins Rib72a-mCherry and Rib72b-mCherry were made using the p4T2-1-NEO2-mCherry vector. In brief, two regions of homology were cloned into the vector flanking the NEO2 gene, which encodes resistance to paromomycin. Homology regions consist of 1kb of DNA from the 3’ end of the *RIB72A* or *RIB72B* gene, terminating at the stop codon and 1kb of DNA from the 3’UTR 100bp after the stop codon of *RIB72A* or *RIB72B.* Rib72a-mCherry and Rib72b-mCherry were also generated with resistance to cyclohexamide. In these strains the *NEO2* coding sequence was replaced by the *RPL38A* coding sequence, which confers resistance to cyclohexamide. Native expression of Rib72a-GFP was generated using the same strategy described above with the mCherry coding sequence replaced with the GFP coding sequence. The *RIB72A* null strain was also generated as described above except the upstream homology region was replaced with 1kb of homologous DNA from the 5’UTR of *RIB72A.* Exogenous expression rescue constructs for N-terminal fusion proteins GFP-Rib72a and GFP-Rib72b were made using the pBS-MTT-GFP-gtw plasmid. In brief, cDNA from *RIB72A* or *RIB72B* was generated by RTPCR using SuperScriptII reverse transcriptase (Invitrogen, Grand Rapids, NY). The cDNA of these genes was cloned into the pENTR4 Gateway Entry Vector (Invitrogen) then shuttled into the pBS-MTT-GFP-gtw plasmid through gateway cloning (Invitrogen). This plasmid integrates into the genome at the *RPL29* locus and its MTT promoter allows for inducible control of expression by using CdCl_2_.

The above described constructs were linearized and introduced to wild type *Tetrahymena* cells by biolistic bombardment. Only the Rib72b rescue construct was integrated into a non-wild type strain background, *RIB72B∆* cells) Plasmid DNA integrated directly into the genome through homologous recombination. Successful integration of plasmid DNA was identified by resistance to either paromomycin or cyclohexamide drug.

The *Tetrahymena* strains lacking one or both of the RIB72 homologs were constructed by gene disruptions (initially in the micronucleus) using homologous DNA recombination and biolistic bombardment (Cassidy-Hanley *et al.*, 1997; Hai and Gorovsky, 1997) as previously described in detail (Dave *et al.*, 2009) using mating strains CU428 and B2068 (obtained from the *Tetrahymena* Stock Center, Cornell University). *RIB72A (TTHERM00143690*) was interrupted by insertion of *neo3* cassette (Shang *et al.*, 2002) while *RIB72B (TTHERM_00584850*) was disrupted by *neo4* (Mochizuki, 2008). Single and double knockout homozygous heterokaryons were obtained by crosses as previously described (Dave *et al.*, 2009). Genotyping of heterokaryon progeny during crosses was performed by PCR of genomic DNA of outcross progeny using primers that detect a junction between the targeted gene and *neo*.

The primers used for construction of targeting plasmids and testing for a loss of the targeted sequences are as follows: Amplification of homology arms for knocking out *RIB72A* were 5’-ATTAGGGCCCAGTGATTAAAACACAGACAAA-3’ and 5’- ATTACCGCGGAGGCTCCCATAAATGATTA-3’; 5’ - TTTATATCGATTATTTTAATTGGCAGAGAGT-3’ and 5’- ATTAGAGCTCATTGATAGGATTGTTGTGAT-3’. Amplification of homology arms for knocking out *RIB72B* were 5’-TTTTGGGCCCTAATTGGTGGTCCTACTGG-3’ and 5’- AAAATCGATACCTTTAGGCTATCCAATCT-3’; 5’ -AAACCGCGGAAGCAATCTAAAAGCCATAGT-3’ and 5’-TTTGAGCTCGTATTAAACTTAGCATTGAATC-3’. Primers to test for the loss of targeted *RIB72A* sequence were 5’- TCTAGGAAAGTGTTGTTGAA-3’ and 5’-ATAGTAATTGAGACCACCTT-3’. Primers to test for the loss of targeted *RIB72B* sequence were 5’-TTTGAGCTCGTATTAAACT TAGCATTGAATC-3’ and 5’-ATGTTCCTCCAG TTGTCTAT-3’.

### Fluorescence microscopy

Generation of the fluorescent fusion *Tetrahymena* strains was achieved as previously described in detail (Winey *et al.*, 2012). Briefly, plasmid DNA was introduced to cells by biolistic bombardment and transformation was mediated by homologous recombination (Cassidy-Hanley *et al.*, 1997; Hai and Gorovsky, 1997). Bbc73 was tagged with mCherry at its C-terminus and integrated at its endogenous locus under control of its native promoter. This fluorescent fusion protein was coexpressed with N-terminal GFP fusions of Cen1 (Vonderfecht *et al.*, 2012), ATU1 or Bbc60 were expressed under control of a inducible promoter (Shang *et al.*, 2002). Fluorescence images were acquired at room temperature using an Eclipse Ti inverted microscope (Nikon, Japan) equipped with a CFI Plan Apo VC 60× H numerical aperture 1.4 objective (Nikon, Japan) and a CoolSNAP HQ2 charge-coupled device camera (Photometrics, Tuscon, AZ). MetaMorph Imaging software (Molecular Devices, Sunnyvale, CA) was used to collect images and generate the projection images. For live cell imaging, cells were grown in 2% SPP media, washed once with 10 mM Tris-HCl, pH 7.4, pelleted, and placed on microscope slides (VWR, Radnor, PA). In strains where gene expression was under control of an inducible MTT1 promotor, 0.5 μg/mL concentration of CdCl2 was used. For immunofluorescence, cells were fixed using either 3% formaldehyde or 70% ethanol, and then placed on poly-L-lysine treated slides. Both the centrin-1 antibody 20H5 and anti-mouse Alexa488 were diluted 1:1000 in PBS+1% BSA. Primary antibody incubation was done at 4°C overnight and secondary antibody incubation was done for 1 hr at room temperature. Cells were washed in PBS+0.1% BSA five times after each antibody incubation. Finally, cells were mounted in Citifluor (Ted Pella, Inc., Redding, CA) and imaged.

### Swim speed assay

*Tetrahymena* cells were grown at 30°C in SSP media (2% protease peptone, 0.1% yeast extract, 0.2% glucose, 0.003% FeCl_3_) to mid-log phase ~2 × 10^5^ cells/mL. Cells were then placed on a slide and imaged using DIC imaging with a 20x objective at 10 frames/sec on a Nikon Ti microscope (Nikon, Japan) using NIS Elements software (Nikon). ImageJ with the ImageJ software plugin MTrackJ (E. Meijering) was used to track and quantify the movement of *Tetrahymena* cells. Twenty cells were quantified per condition, and the experiment was performed in triplicates.

### Reciliation assay

*Tetrahymena* cells were grown in 2% SPP media to mid-log phase ~2 × 10^5^ cells/mL. Deciliation was performed as described previously(Calzone and Gorovsky, 1982). Briefly, cells were placed in starvation media (10 mM Tris-HCl, pH 7.4) for 24 hours at 30°C. Cells were then placed in deciliation medium (10% Ficoll 400, 10mM NaOAc, 10mM EDTA, 10 mM CaCl_2_, pH 4.2) and cilia were mechanically sheared by passage through an 18.5 gauge syringe. Cells were then placed in regeneration medium (15 mM Tris-HCl, 2.0 mM CaCl_2_ pH 7.95) and allowed to regenerate cilia at 30°C. Cells were imaged every thirty minutes as described above in the swim speed assay. Twenty cells from each time point were scored for forward movement (>25 μm/sec) and the percentage of the 20 cells that showed forward motility was recorded. The experiment was completed in triplicates.

### Phagocytosis assay

*Tetrahymena* cells were incubated with 0.1% India ink in 2% SSP media at 30°C. At 0.5, 2, 4.5, and 7.5 h after the India ink was added, cells were washed in 10 mM Tris buffer (pH 7.5) and fixed with 4% glutaraldehyde (Sigma-Aldrich). Cells were mounted on slides and the average number of food vacuoles per cell was quantified (n=10 cells) using phase contrast imaging with a 60x oil immersion objective on an upright light microscope (Nikon Eclipse LV, Japan) equipped with sCMOS camera (Andor, UK).

### High speed video analysis of cilia in swimming *Tetrahymena* cells

Observations of cilia in actively-swimming *Tetrahymena* cells were performed and data were quantified as described previously for *Paramecium tetraurelia* (Funfak *et al.*, 2015), with minor modifications. A highly concentrated cell suspension (in 2% SPP medium) was injected into the inlet of a microfluidic channel (33 μm in height and 50 μm in width) using a controllable pressure source (Fluigent). The pressures between the inlet and outlet of the channel were equilibrated to prevent residual flow. Differential interference contrast (DIC) imaging of forward-swimming cells was done on an inverted microscope (Nikon TE2000), using a 60x objective lens. Videos were recorded on a Photron 1024PCI camera at 2000 frames per second. Video image processing and quantification was done exactly as described (Funfak *et al.*, 2015). Matlab (The MathWorks Inc., Natick, MA) along with the CR Toolbox (Adelin Barbacci, 2014) were used to mark the cell body and the zone of beating cilia in the first frame of the movie. The cell body and ciliary zone boundaries was adjusted every 30-50 frames to track the position of the moving cell. The segment of the video containing cilia was digitally unwrapped, producing a rectangle where all ciliary bases are aligned horizontally, generating an “extracted cilia video”. ImageJ was used to plot the grey-scale over time at a line positioned within the ciliary zone but away from the cell surface, creating a space-time diagram (chronograph). This produces an image containing a number of short diagonal lines that correspond to traces of individual cilia. On the time (y) axis, successive diagonal lines corresponding to the same cilium can be identified based on their vertical position and shape. The beat frequency was quantified by measuring the distance (on the y-axis) between diagonal lines corresponding to the same cilium. The speed of the power stroke corresponded to the inverse slope of a diagonal line. See Figure S2 for additional information about the methodology used to quantify ciliary motility.

### Cryo-sample preparation and cryo-ET

Cryo-samples were prepared and imaged as previously described (Nicastro *et al.*, 2006; Nicastro *et al.*, 2011). Briefly, axonemes were isolated from cultured *Tetrahymena* cells using the pH-shock method. When the pH of the buffer solution was rapidly lowered to 4.3, the cells released their intact cilia, which were collected by centrifugation. Cilia were then demembranated with detergent (IGPAL 630, Sigma-Aldrich) and the axonemes collected by centrifugation. 10 nm colloidal gold solution (Sigma-Aldrich) was applied to Quantifoil grids R2/2 Holey carbon 400 mesh Quantifoil grids (Quantifoil MicroTools GmbH, Germany) and allowed to air-dry. 3 μL of sample and 1 μL of 10-fold concentrated BSA-coated 10-nm gold solution (Iancu, 2006) were added to the grid, and excess fluid was blotted with filter paper before the grid was rapidly plunge frozen in liquid ethane with a homemade manual plunge freezing device. This generates a layer of vitreous ice with embedded axonemes. Frozen grids were stored in liquid nitrogen until used.

For cryo-ET, frozen grids were loaded into a cryo-holder (Gatan, Inc. Eindhoven, Netherlands) and inserted into a Tecnai F30 transmission electron microscope (FEI, Inc. Hillsboro, OR) equipped with a field emission gun, GIF2000 post column energy filter (Gatan, Inc. Eindhoven, Netherlands), and a 2,048 × 2,048 pixel CCD camera (Gatan, Inc. Eindhoven, Netherlands). Single-axis tilt series were recorded using the microscope control software SerialEM (Mastronarde, 2005) with the TEM operated at 300 kV in low-dose mode. The sample was tilted from −65 to +65° in increments of 2.0°. The total accumulated electron dose was limited to ∼100 e/Å^2^ The defocus was −8 μm, and the energy filter was operated in zero loss mode with a 20 eV slit width. The magnification was 13,000x with a final pixel size of 1.077 nm.

### Image Processing

The software program IMOD (Kremer *et al.*, 1996) was used for fiducial alignment and tomogram reconstruction using weighted back projection. Subtomogram averaging was performed using the PEET software (Nicastro *et al.*, 2006; Heumann *et al.*, 2011) to average the 96-nm axonemal repeat units from 3D reconstructed *Tetrahymena* axonemes. The resolution of the DMT averages was estimated using the 0.5 criterion of the Fourier shell correlation method (supplementary Table 1). The Chimera package (Pettersen *et al.*, 2004) was used for 3D visualization and surface rendering

### Online Supplemental Material

Fig. S1 shows fluorescently tagged RIB72A and RIB72B localizing to cilia and basal bodies independently. Stained phagocytic vacuoles show phagocytosis defects in Rib72BKO and Rib72A/BKO. Overexpression of RIB72B-GFP rescues swim speed phenotype in Rib72BKO cells. Fig. S2 depicts drawings that explain the generation of chronographs from high speed video images of moving *Tetrahymena.* Fig. S3 shows significantly decreased power stroke speed for mutant cilia. Also shows high speed video frames tracing the power stroke of WT and mutant cilia. The 3D isosurface renderings of the axonemal repeats of WT and mutants in Fig. S4 demonstrate that the external microtubule-associated complexes are not defective in the here studied mutants. Table S1 shows strains used in this study, along with image processing information where applicable. Video S1-S4 show *Tetrahymena* cells swimming inside a microfluidic channel, recorded at 2000 frames/sec from wild-type (video S1), RIB72A-KO (video S2), RIB72B-KO (video S3) and RIB72A/B-KO (video S4), respectively. Video S5 shows a RIB72A/B-KO cell swimming inside a microfluidic channel, recorded at 2000 frames/sec. Red arrow marks a cilium with a kink near the distal region.

## Acknowledgements

We thank Chen Xu for training and management of the electron microscopes in the Louise Mashal Gabbay Cellular Visualization Facility at Brandeis University (Waltham, MA). This work was supported by funding from the National Institutes of Health (GM083122 to DN, GM074746 and GM127571 to MW, and GM089912 to JG) and the National Science Foundation (MBC-033965 to JG). PL was supported by a predoctoral fellowship from the American Heart Association. The authors declare no competing financial interests. The 3D averaged structures have been deposited in the Electron Microscopy Data Bank (EMDB) under accession code EMD-7806, EMD-7811, EMD-7805 and EMD-7807. Author contribution: D.N., M.W. and J.G. conceived and planned experiments. D.S. carried out sample preparation, cryo-electron microscopy, subtomogram averaging and phagocytosis experiments. D.D., J.G., S.S., and Y.Z. generated *Tetrahymena* mutants and performed genetic experiments. Y.Z. also performed fluorescent microscopy. B.A.B. performed phenotypical characterization of *Tetrahymena* strains. P.L., R.F.-X.T., P.D.-W. and C.N.B. performed the high speed video imaging and waveform analysis. All authors contributed to data analysis and interpretation. D.S., L.G. and D.N. wrote the manuscript with input from all authors.

## Abbreviations

A_t_: A tubule
B_t_: B tubule
cryo-ET: cryo-electron tomography
DMT: doublet microtubule
IDA: inner dynein arms
MIP: microtubule inner protein
N-DRC: nexin-dynein regulatory complex
ODA: outer dynein arms

---

**Figure S1.**
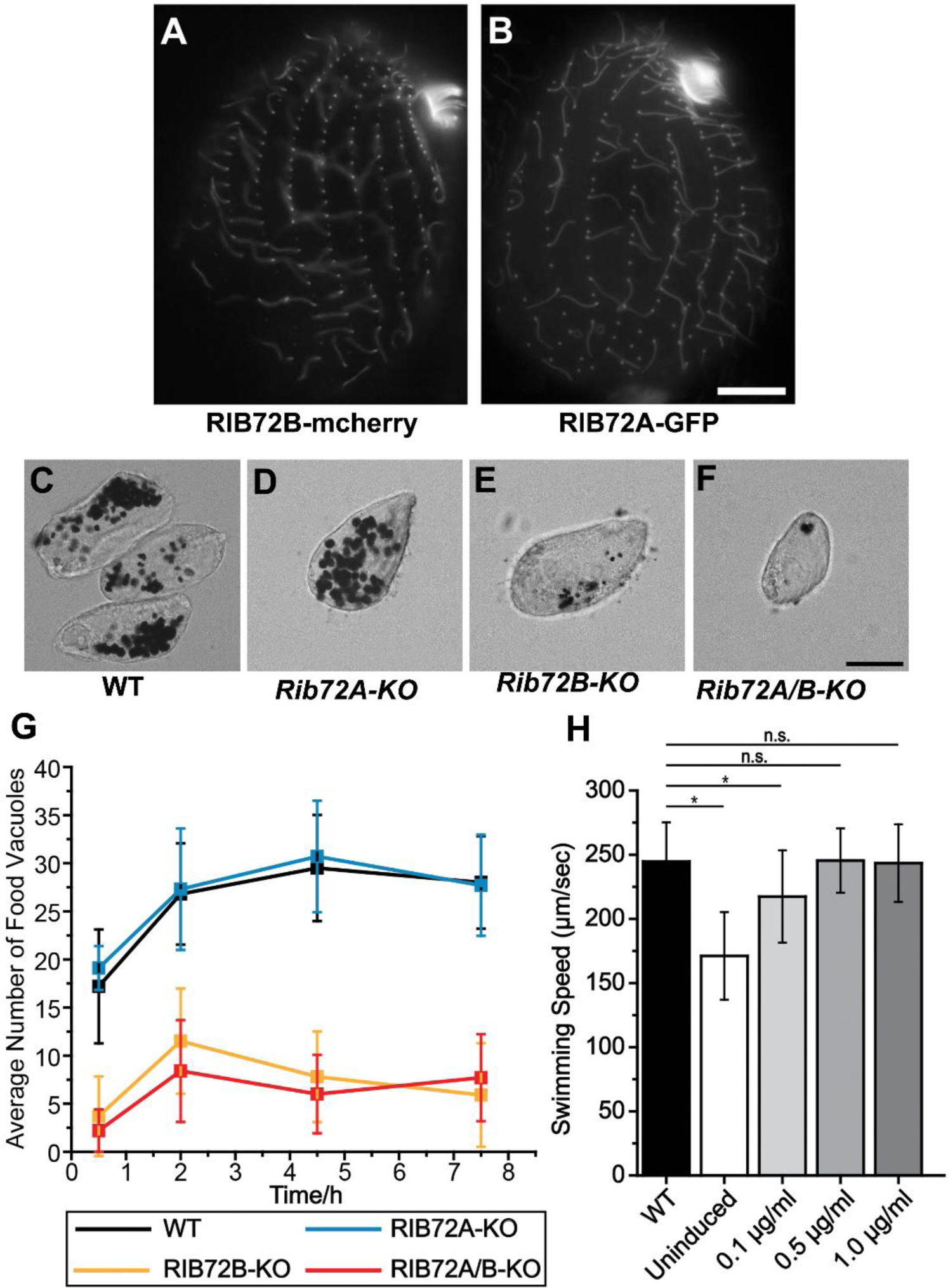
RIB72A and RIB72B localize independently of each other to cilia and basal bodies. (A, B) Fluorescent light microscopy images of fixed *Tetrahymena* cells shows (A) GFP-tagged RIB72A in the *RIB72B-KO* mutant background, and (B) RIB72B-mCherry in the *RIB72A-KO* mutant background still localizing to cilia and basal bodies. Scale bars: 10 μm. **(C-F)** Phase contrast images of fixed *Tetrahymena* cells from WT (C), *RIB72A-KO* (D), *RIB72B-KO* (E) and *RIB72A/B-KO* (F) showing phagocytic vacuoles stained with India ink that was provided in the growth medium. Scale bar: 20 μm. **(G)** The number of phagocytic vacuoles was recorded at different time points after addition of the India ink. The most severe phagocytosis defects were evident in *RIB72B-KO* (E) and *RIB72A/B-KO* (F). (H) The expression level of RIB72B-GFP is induced by different levels of cadmium. Error bars represent standard deviations and asterisks indicate statistical significance at p<0.01 (one way ANOVA); n.s., non-significant, p>0.05.

**Figure S2.**
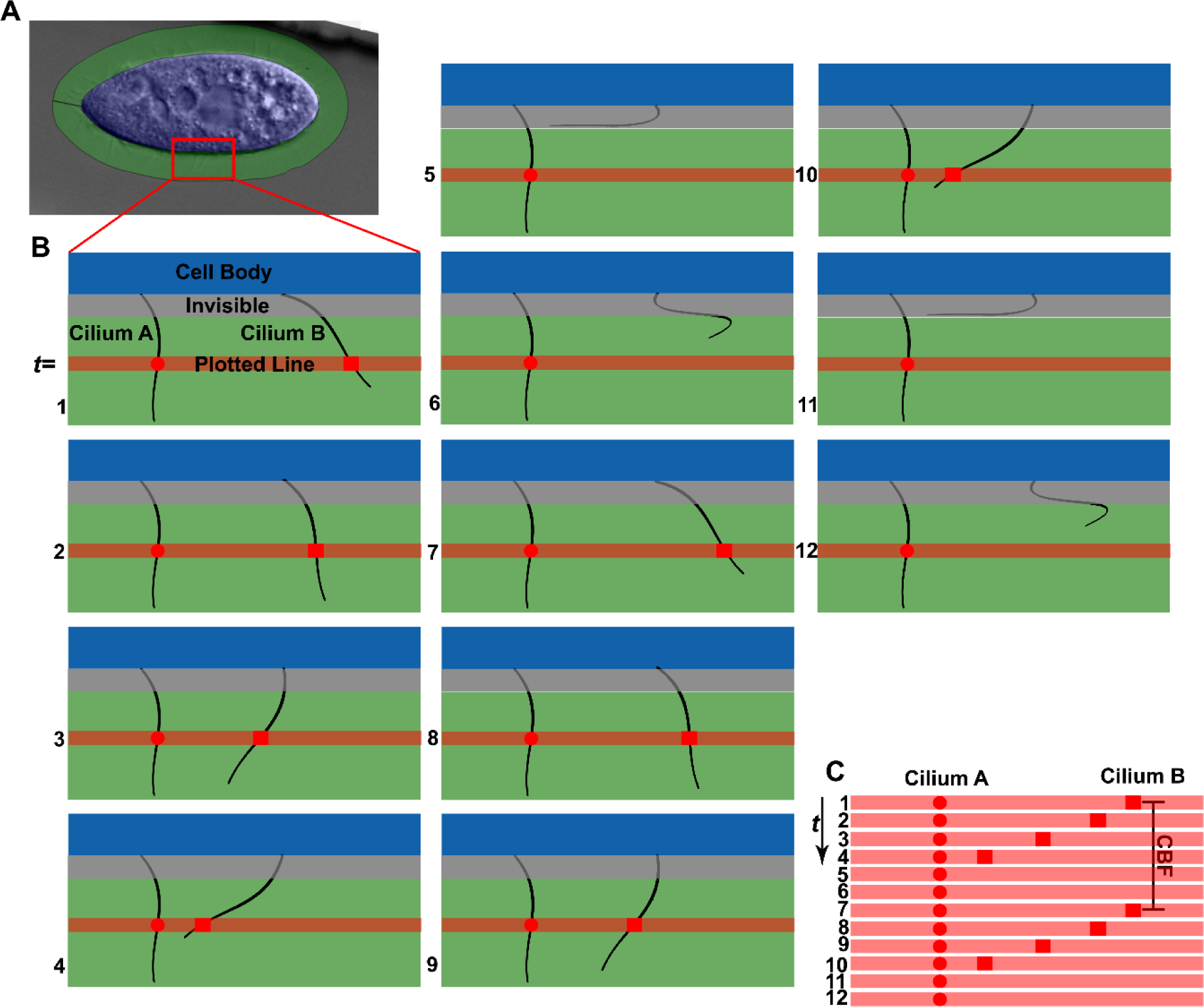
Schematic drawings explaining the generation of chronographs based on highspeed video imaging (shown in Fig. 4). (A) First frame of a video was used to mark the cell body (blue) and cilia regions (green) of a swimming *Tetrahymena* cell. The red box outlines an area in the cell periphery that is shown at higher zoom in (B). **(B)** Selected area of the cell periphery of a swimming *Tetrahymena* cell at different time-points t=1–12 in chronological order. Two cilia are drawn: cilium A is stalled, whereas cilium B undergoes two complete beating cycles between t=1 and t=12. Note that beating cilia are not visible during their recovery stroke (e.g. cilium B at time- points 5,6 and 11,12), because the cilium is moving too close to the cell body and “disappears” in the fringe close to the cell caused by DIC imaging. The red lines show the position which was plotted over time to produce the chronograph seen in (C); note the red circle and squares on the red line that indicate where the line transects a cilium. **(C)** Chronograph of cilia A and B constructed from lines that transect the video frames at the same location over time. Cilium A is stalled and does not change position throughout the Chronograph as indicated by the vertical column of circles. Cilium B is actively beating and changes position over time t as indicated by the red squares. The cilia beat frequency (CBF) is the time required for one complete beating cycle including the recovery stroke.

**Figure S3.**
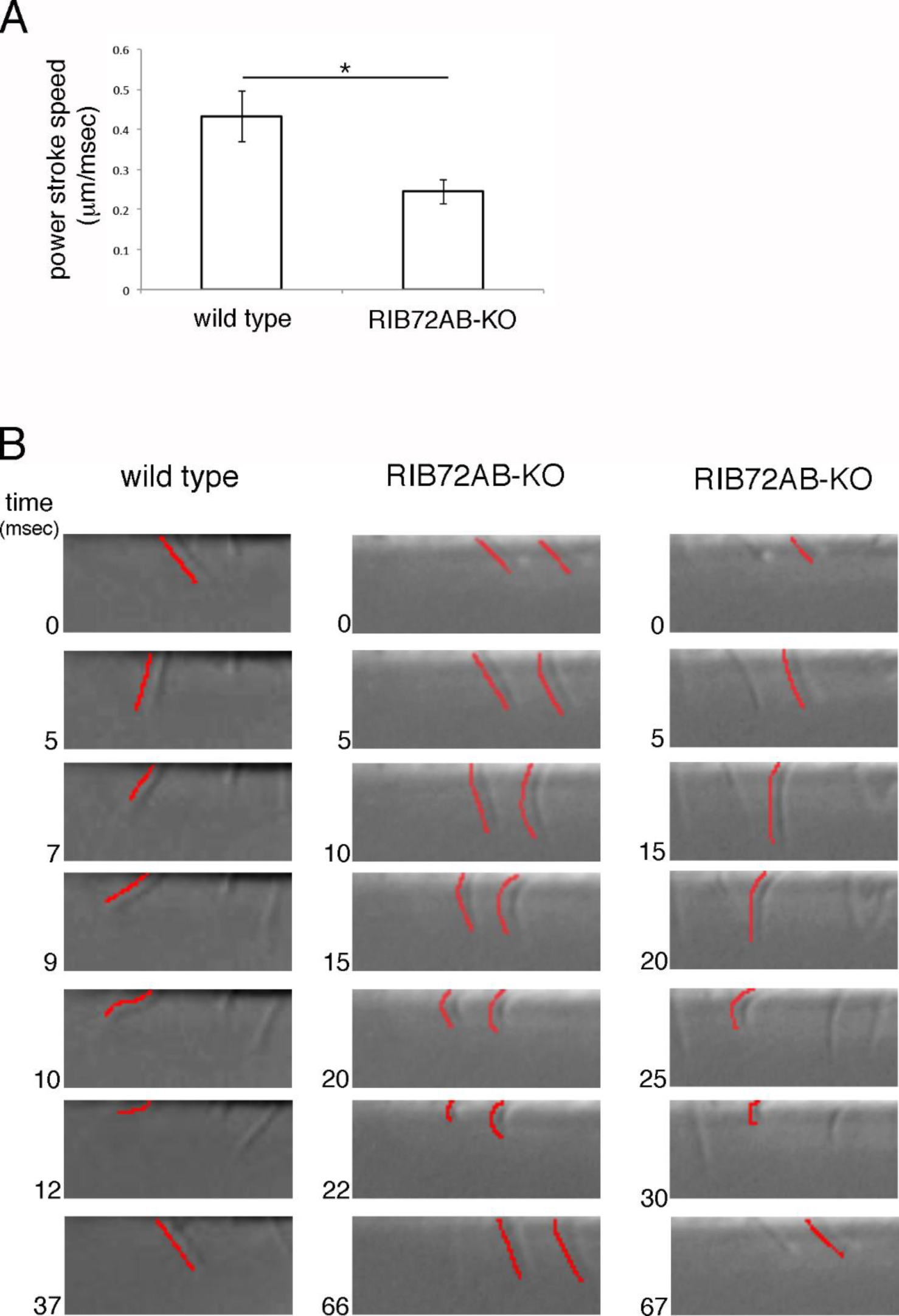
*RIB72A/B-KO* cilia have a slower power stroke and waveform defects. (A) A graph showing the power stroke speed values in wild-type (0.43 μm/msec, n cilia = 10, n cells =3) and *RIB72A/B-KO* (0.24 μm/msec n cilia = 10, n cells =3) cilia. The values were extracted from the chronographs (Fig. 1B-E); the inverse slope of a short diagonal line of a cilium trace represents the speed of the power stroke. Error bars represent standard deviation and asterisk shows statistical significance at p< 0.001 (t-test). **(B)** Successive frames showing examples of individual cilia obtained from wild-type and *RIB72A/B-KO* extracted cilia videos (video S1, video S4). The red line in each frame traces the shape of the same cilium. Note that cilia are invisible in most of the recovery stroke as their shafts lie close to the cell surface.

**Figure S4.**
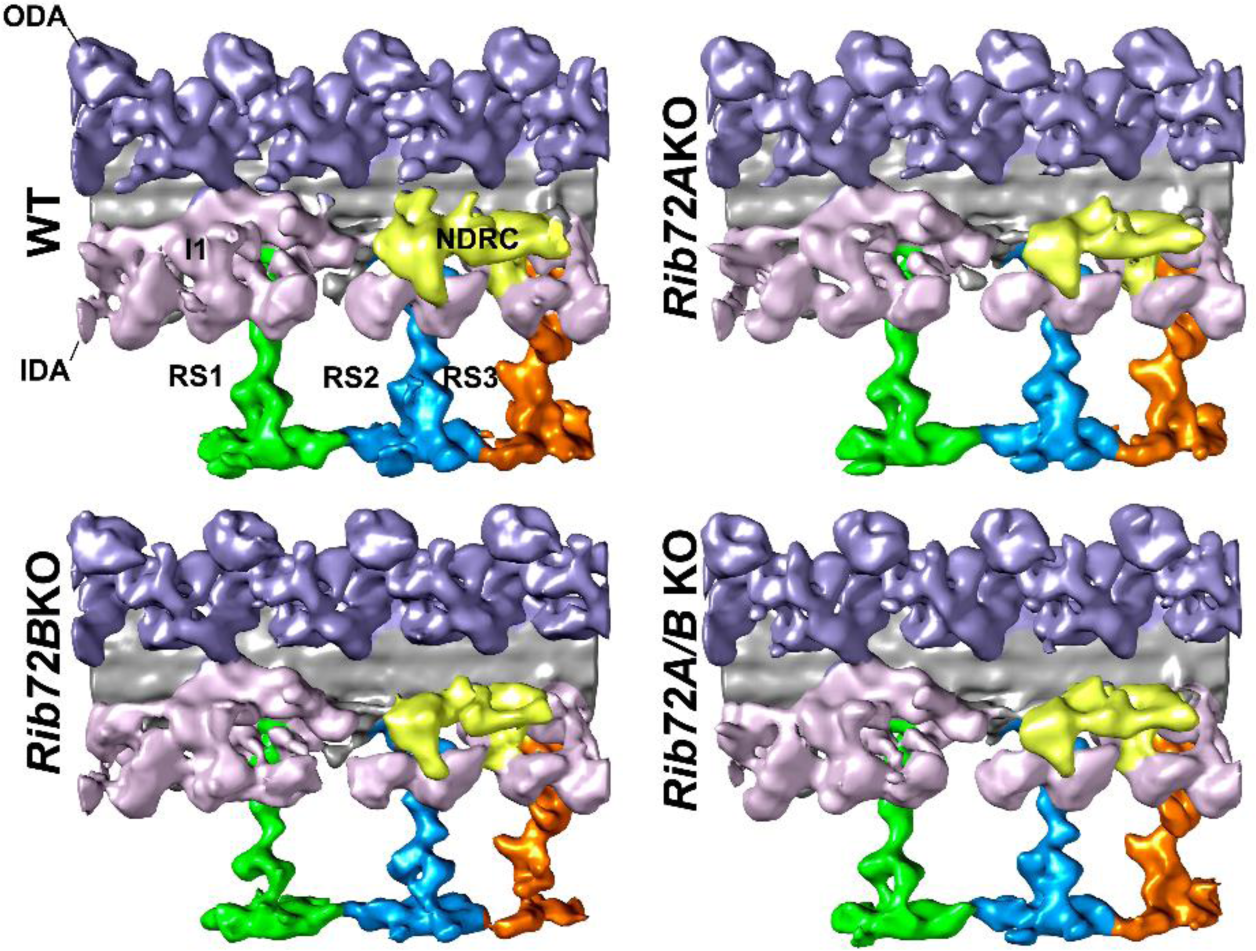
All axonemal structures associated to the outside of the DMTs appeared unchanged in the mutants. (A-D) Isosurface renderings in longitudinal front view (proximal on the left) of *Tetrahymena* axonemal repeat units of WT (A), *RIB72A-KO* (B), *RIB72B-KO* (C), and the double mutant (D). Colors: Inner dynein arms IDA (rose); nexin dynein regulatory complex N-DRC (yellow); outer dynein arms ODA (lavender), radial spokes 1 (green) 2 (blue) 3 (orange).

**Supplementary Table 1.** Strains used in this study and image processing information where applicable

| Name | Strain | LM only | Cryo-ET:<br># of tomograms included | Averaged repeats | Resolution<sup>a</sup> (nm) | Used in figure(s) |
| --- | --- | --- | --- | --- | --- | --- |
| WT<sup>b</sup> | CU428 |  | 7 | 500 | 3.4 | 3-8, S1,S3,S4 |
| *RIB72A-KO* | *RIB72A-KO* |  | 5 | 758 | 4.4 | 3-7<sup>c</sup>, S1<sup>d</sup>,S4 |
| *RIB72B-KO* | *RIB72B-KO* |  | 5 | 702 | 4.4 | 3-7, S1,S4 |
| *RIB72A/B-KO* | *RIB72A/B-KO* |  | 5 | 699 | 4.4 | 3-7 S1,S3,S4 |
| *RIB72B-GFP* | *RIB72B-KO:*<br>*RIB72B-GFP* |  | 5 | 735 | 4.6 | 8 |
| RIB72A-mCherry, GFP-CEN1 | RIB72A-mCherry, GFP-CEN1 | X |  |  |  | 2 |
| RIB72A-mCherry, GFP-ATU | RIB72A-mCherry, GFP-ATU | X |  |  |  | 2 |
| RIB72B-mCherry, GFP-RIB72A | RIB72B-mCherry, GFP-RIB72A | X |  |  |  | 2 |
| *GFP-RIB72A* | *RIB72B-KO:*<br>*GFP-RIB72A* | X |  |  |  | S1 |
| *RIB72B-mCherry* | *RIB72A-KO:*<br>*RIB72B-mCherry* | X |  |  |  | S1 |

<sup>a</sup>The 0.5 Fourier shell correlation criterion was used to estimate the resolution of the averaged repeats at the site of the doublet microtubule.
<sup>b</sup>WT includes previous published data (Vasudevan, Song et al. 2015).
<sup>c</sup> *RIB72A-KO* generated by Winey Lab
<sup>d</sup> *RIB72A-KO* generated by Gaertig lab

## Supplemental Video Legends

**Video S1.** A wild-type *Tetrahymena* cell swimming inside a microfluidic channel, recorded at 2000 frames/sec. (Stills appear in Figure 4 and Figure S3)

**Video S2**. A *Tetrahymena RIB72A-KO* cell swimming inside a microfluidic channel, recorded at 2000 frames/sec. (Stills appear in Figure 4)

**Video S3.** A *Tetrahymena RIB72B-KO* cell swimming inside a microfluidic channel, recorded at 2000 frames/sec. (Stills appear in Figure 4)

**Video S4.** A *Tetrahymena RIB72A/B-KO* cell swimming inside a microfluidic channel, recorded at 2000 frames/sec. (Stills appear in Figure 4 and Figure S3)

**Video S5.** A *Tetrahymena RIB72A/B-KO* cell swimming inside a microfluidic channel, recorded at 2000 frames/sec. Red arrow marks a cilium with a kink near the distal region.

## Condensed title

RIB72A and RIB72B are Microtubule Inner Proteins

